# Analysis of Fox genes in *Schmidtea mediterranea* reveals new families and a conserved role of *Smed-foxO* in controlling cell death

**DOI:** 10.1101/2020.09.11.292938

**Authors:** Eudald Pascual-Carreras, Carlos Herrera-Úbeda, Maria Rosselló, Jordi Garcia-Fernandez, Emili Saló, Teresa Adell

## Abstract

The forkhead box (Fox) genes encode transcription factors that control several key aspects of development. Present in the ancestor of all eukaryotes, Fox genes underwent several duplications followed by loss and diversification events that gave rise to the current 25 families. However, few Fox members have been identified from the Lophotrochozoa clade, and specifically from planarians, which are a unique model for understanding development, due to the striking plasticity of the adult. The aim of this study was to identify and perform evolutionary and functional studies of the Fox genes of lophotrochozoan species and, specifically, of the planarian *Schmidtea mediterranea*. Generating a pipeline for identifying Forkhead domains and using phylogenetics allowed us the phylogenetic reconstruction of Fox genes. We corrected the annotation for misannotated genes and uncover a new family, the QD, present in all metazoans. According to the new phylogeny, the 27 Fox genes found in *Schmidtea mediterranea* were classified into 12 families. In Platyhelminthes, family losses were accompanied by extensive gene diversification and the appearance of specific families, the A(P) and N(P). Among the newly identified planarian Fox genes, we found a single copy of *foxO*, which shows an evolutionary conserved role in controlling cell death.

**Author summary:** Transcription factors are the key elements that regulate gene expression in the nucleus. The forkhead box (Fox) transcription factors are one of the most numerous and they control key aspects of development. Fox genes were already present in the ancestor of all eukaryotes, and then underwent several duplications followed by loss and diversification events that gave rise to the current Fox families in the different species. The available data classifies Fox genes in 25 families, but they include few members corresponding to Lophotrocozoa, one of the two invertebrate phyla that includes annelids, molluscs or platyhelmintes. In this study we identify and perform evolutionary studies of the Fox genes of several lophotrochozoan species and, specifically, of the planarian *Schmidtea mediterranea*. The result is the correction of the annotation of Fox genes from many species, proposing a new nomenclature, and the identification of new families; the QD family, present in all metazoans, and the A(P) and N(P) families, specific of Platyhelminthes. We also study the function of *Schmidtea mediterranea foxO*, a gene involved in aging and cancer in other species, showing its evolutionary conserved role in controlling cell death according to cell metabolism.

## Introduction

Forkhead box (Fox) genes belong to the ‘winged helix’ superfamily of transcription factors (TF) with a specific DNA-binding domain referred to as the Forkhead domain (FKH), with approximately 100 Aa. In Metazoa, Fox genes are expressed in a specific spaciotemporal manner during development, and control essential processes as cell death, cell cycle and stem cell differentiation into specific cell linages and populations (1–3). Thus, Fox genes play a major role during embryonic and postembryonic development, such as: gastrulation, lifespan, immune system regulation or tissue differentiation and maintenance (1). In humans, the lack of some Fox genes leads to embryonic lethality or developmental diseases such as Parkinson’s, defects in the immune system, speech and language learning or cancer (1,2,4,5).

Currently over 2000 Fox proteins have been identified in 108 species of fungi and metazoans, including a vast number of Phyla, such as Choanoflagellata (6), Ctenophora (7), Placozoa (8), Porifera (9), Cnidaria (10), Echinodermata (11), Hemichordata (12), Cephalochordata (13,14) and Chordata (14,15). However, few Fox genes have been identified in lophotrochozoan species, and most functional studies have been only performed in few model organisms such as mice (16–18), *Drosophila* (19,20) or *C. elegans* (21). Planarians are Lophotrochozoans well known for their astounding ability to regenerate any body part and change their size according to food availability. Such tissue plasticity is due to the presence of adult stem cells (neoblasts) that can give rise to all differentiated cell types, which is accompanied by the continuous activation of the intercellular signalling mechanisms (22–24). Planarians’ phylogenetic position and plasticity makes them an interesting model for investigating the Fox family at an evolutionary and functional level.

Fox genes are currently phylogenetically classified and grouped into 25 families (A to S) (2,25). The different gains and losses of Fox families have shaped the history of Fox family evolution, such as the division of family Q into Q1 and Q2, N into N1/4 and N2/3, L into L1 and L2 or J into J1 and J2/3 (14). Another example of gain is the S family, which seems to evolve by duplication of the C family and is specifically found in vertebrates (11,26). Fox family losses have also been reported, such as the AB in vertebrates (15) or E, H, I, J2/3, M and Q1 in Ecdysozoa (14).

The aim of this study was to identify and classify the Fox genes of the planarian species *Schmidtea mediterranea* (*Smed*), along with the Fox genes of other lophotrochozoan species whose genome or transcriptome is currently available (27–32). Previous studies of Fox genes expression and function in *Smed* showed that they were tissue specific and participated in its maintenance. The essential role of *foxA* in the maintenance of the pharynx and endodermal tissue has already been identified in planarians (33), similar to its role in early endoderm development in vertebrates (34–36), or the role of J1 paralogs in ciliogenesis (37), similar to the one described in vertebrate species as mice, chickens or frogs (38,39). However, other important Fox families such as the O family, related with metabolism, growth and aging (40,41) had not been identified in planarias.

Through generating a pipeline for identifying Forkhead domains we identified and annotated 27 Fox genes in *Smed*, 18 of which are firstly reported here. Phylogenetic analyses allowed us to classify *Smed* Fox in 12 families. The integration of the newly identified Fox from *Smed* and other lophotrochozoans and Platyhelminthes with all reported Fox genes allowed for the identification of the QD family, a new family which appears to originate after the split of sponges from the rest of the eumetazoans. Most of the Fox families also originated during this period, which was followed by various loss events and some diversification. Specifically, Platyhelminthes suffered a huge gene family loss followed by gene diversification originating specific families: A(P) and N(P). Finally, we identified a single copy of the *foxO* gene in planarians and demonstrated its conserved role in controlling cell death.

## Results

### *Schmidtea mediterranea* presents 27 Fox genes that can be classified in 12 families

With the aim to identify all Fox genes of *Smed*, we developed a pipeline for identifying Forkhead domains (FKH) using the available FKH from Pfam in combination with TransDecoder and HMMER (Fig. 1a, see Methods). As a result, we found 27 distinct genes that contained this domain. To determine which family each of these genes belonged to, we performed a phylogenetic analysis using the FKH domain of the Fox genes of an additional 20 species across metazoans, including several lophotrochozoans, to better resolve the *Smed* Fox groups (Fig. 1a and methods). The analysis resulted in the classification of the 27 FKH-containing *Smed* genes into 12 Fox families (Fig. 1b and c, S1, S2, S3). The complete information and new annotation regarding each FKH-containing gene identified is provided in the supplementary materials (S1 File), along with the raw tree (S2 File).

**Figure 1.**
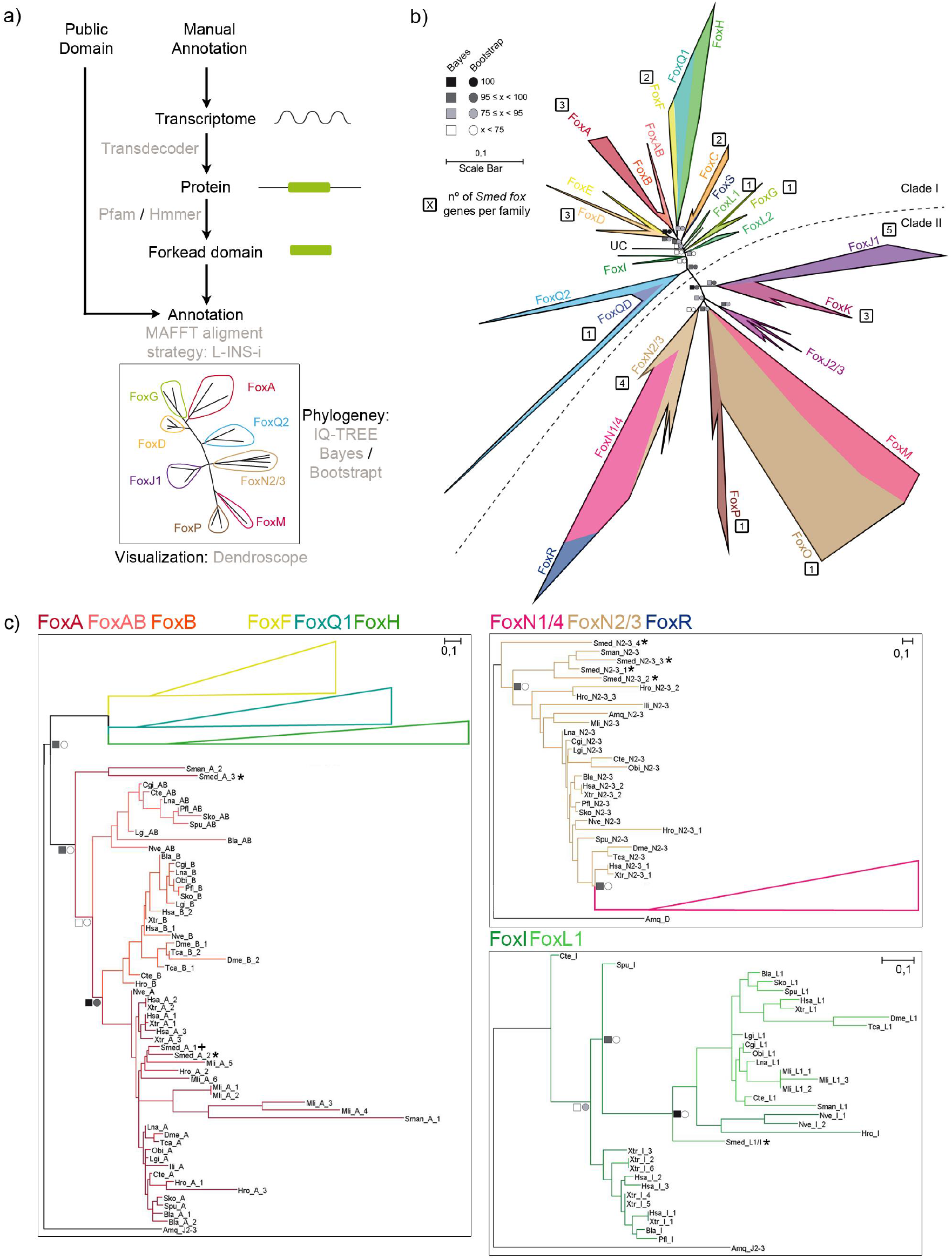
Fox family evolution in Metazoa reveals 27 Fox genes in *Schmidtea mediterranea* divided in 12 families. **a** pipeline annotate Fox genes. **b** The ML phylogenetic trees based on FKH. Number of genes per family in *Schmidtea mediterranea* is indicated inside a square next to each family. At nodes, values for the approximate Bayes (square) and Likelihood (circle) ratio test are shown. Colour indicates % of confidence. Family tree branches were collapse at the base of common node. One gene was unclassified in any family (UC). Dashed line divides Clade I and Clade II Fox genes. **c** For each node-sharing families, a phylogenetic tree was created using an *Amq* gene from the opposite clade as out group. Family branches are painted with the same colour as they are represented in the trees. Dark crosses indicate previous characterized genes and dark asterisks indicate new fox characterized in *Schmidtea mediterranea* (*Smed*). Aminoacidic sequences used are found in S1 File. Scale indicates expected aminoacidic substitution per site.

The phylogenetic tree (Fig. 1b) shows that 5 out of these 12 families belong to Clade II, which is argued to be the ancestor clade (42,43); and 7 belong to Clade I. To better visualise the different Fox genes in each cluster, we inferred a series of new phylogenetic trees including only the genes from closely related families (Fig. 1c, S2, S3). Using this visualisation, we found some *Smed* Fox genes that were not properly classified: a *Smed* Fox gene that clusters between the L1 and the I families (*Smed-foxL1/I*) (Fig. 1c), a *Smed* Fox gene clustering as a sister group of the A, AB and B families (*Smed-foxA3*) (Fig. 1c, S2), and a *Smed* Fox clustering as a sister group of the N2/3 family (*Smed-foxN2/3-4*) (Fig. 1c, S3). Furthermore, we can observe how the Q2 family, widely described in many species (12,44–46) has a branch populated with several genes that cluster with a divergent Q2 gene known as *foxQD* in *Saccoglossus kowalevskii* (*Sko*) (12). We consider this branch to be a new family (Fig. S2) which we called QD, due to the FKH-containing gene originally described in *Sko*. A *Smed* Fox gene belongs to this family (*Smed-foxQD*).

Focusing on the presence of Fox genes in each family, we can observe that despite the number of Fox genes in *Smed* has remained similar to the rest of lophotrochozoans (see purple lines in Fig. 2), the number of families with representatives of *Smed* and the other two Platyhelminthes (*Schistosoma mansoni* and *Macrostumum lignano*) has decreased. Particularly, Platyhelminthes seem to have lost the AB, B, E, H, I, Q1, Q2, M and N1/4 families (red dashed square in Fig. 2). This suggests a huge family loss at the base of Platyhelminthes phylum (orange lines in Fig.2) coupled with an expansion of the Fox number in specific families.

**Figure 2.**
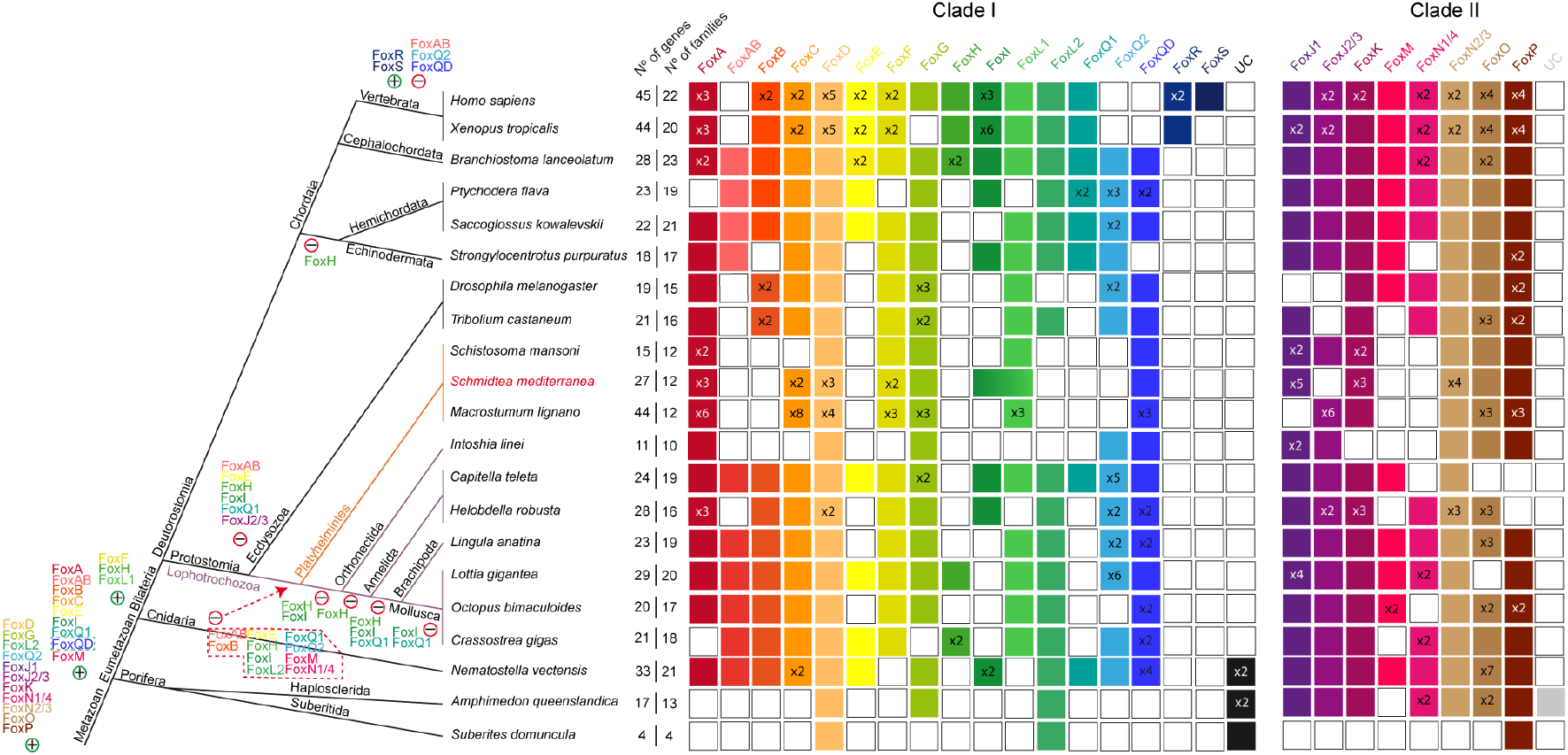
Distribution of Fox homologs in Metazoan clade indicating gene and family losses and some gene duplications in *Schmidtea mediterranea*. Coloured boxes indicate the presence of an ortholog based on the phylogenetic analysis. When there were no evidences of ortholog, box remains white. A number (x N°) inside a box indicates paralogs per family and specie. Families are divided in Clade I and II. Number of genes and number of families per species are indicated. Metazoan (103) and Lophotrochozoa (104) phylogenies were used. Light purple lines indicate lophotrochozoan species and within light orange indicates Platyhelminthes superphylum species. Gains (+) and a losses (−) of genes are placed next to each clade. Main Clade I Fox acquisition was at base of Eumetazoa and different events of gains and losses happened thought evolution. Specifically, many families were lost in Platyhelminthes (red dashed polygon).

### Platyhelminthes present specific Fox subfamilies: FoxA(P) and FoxN(P)

To further investigate the phylogeny of the unclassified *Smed* Fox genes (*Smed-foxL1/I*, *Smed-foxA3* and *Smed-foxN2/3-4*) we performed a second phylogenetic analysis focused on Platyhelminthes data. Repeating the same pipeline previously described, we identified the FKH-containing genes from a total of 19 Platyhelminthes species (including *Smed*), 8 of which belong to the Tricladida order, to which *Smed* belongs (Fig. 3a). Platyhelminthes Fox data can be found in the S3 File and the raw tree can be found in the S4 File. As previously, we also performed additional phylogenetic trees of close-related families to better visualize each family (Fig. S1, 3b, S4, S5). This analysis allowed us to properly classify the FoxL1/I gene into the L1 family (Fig. 3b), which seems to be slightly divergent in the Tricladida Order, and thus we renamed it as *Smed-foxL1.* Furthermore, the new analysis allowed the identification of two new subfamilies only present in Platyhelminthes to which the *Smed-foxA3* and the *Smed-foxN2/3-4* genes belonged. Thus, we renamed them as *Smed-foxA(P)* (Fig. 3b) and *Smed-foxN(P)*, respectively (P meaning specific of Platyhelminthes) (Fig. S5). In this analysis the N subfamily was found to be specific of Triclads, while the A(P) subfamily was also found in other Platyhelminth orders.

**Figure 3.**
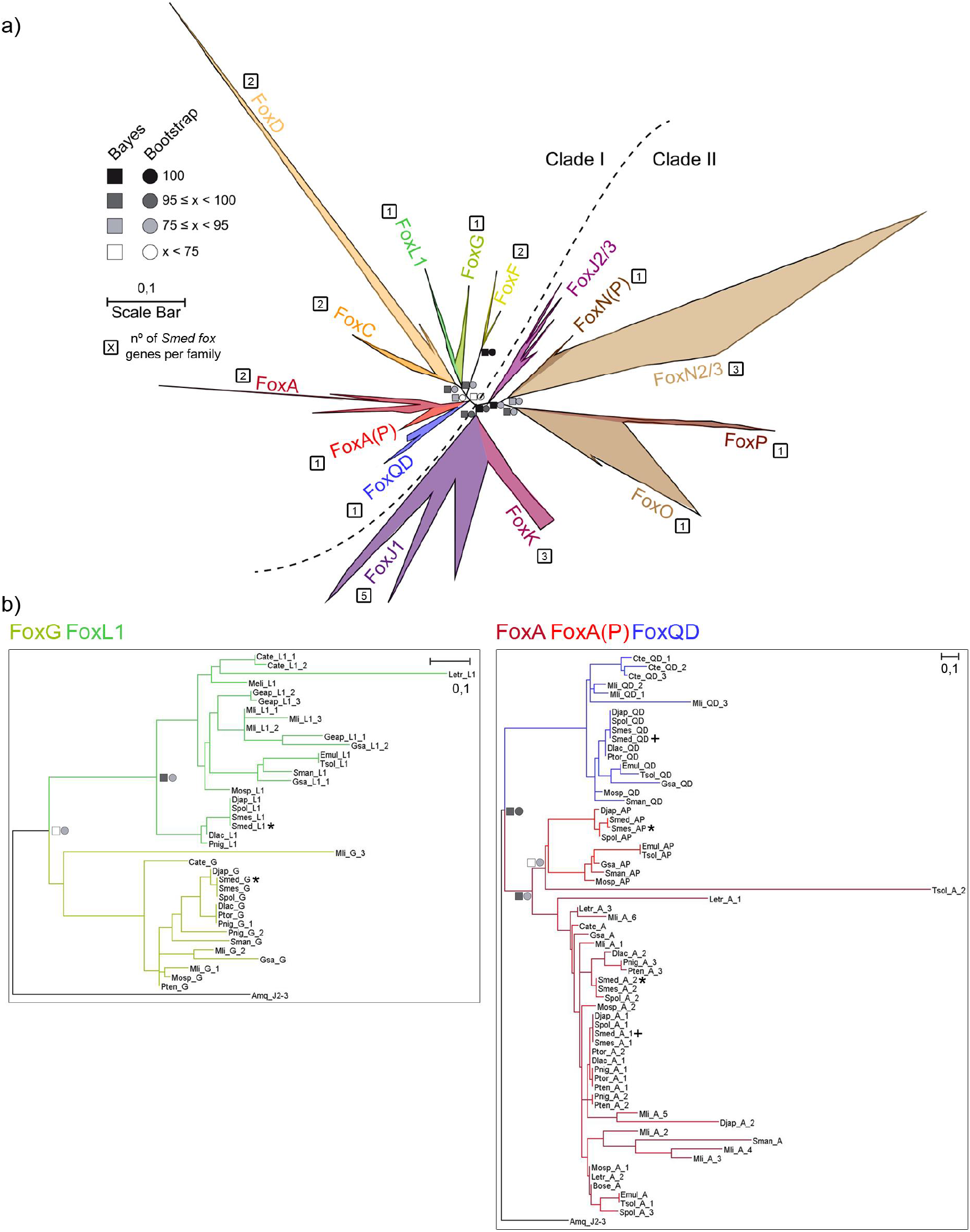
Fox family evolution in Platyhelminthes indicates family diversification. **a** The ML phylogenetic trees based on FKH of Fox family evolution in Lophotrochozoan clade. Number of genes per family in *Schmidtea mediterranea* are indicated inside a square next to each family. At nodes, values for the approximate Bayes (square) and Likelihood (circle) ratio test are showed. Colour indicates % of confidence. Family tree branches were collapse at the base of common node. Dashed line divides Clade I and Clade II Fox genes. **b**For each node-sharing families, a phylogenetic tree was created using *Amq* gene from the opposite clade as out group. Family branches are painted with the same colour as they are represented in the trees. Dark cross indicates previous characterized gene and dark asterisk indicates new fox characterized in *Schmidtea mediterranea* (*Smed*). Aminoacidic sequences used are placed in S3 File. Scale indicates expected aminoacidic substitution per site.

Based on these analyses we have identified and classified all FKH containing genes of *Smed*, including the ones already published, which in some cases have been reclassified according to our analysis. Thus, the previous *Smed-foxQ2* (47) is now classified as *Smed-foxQD*, and the previous *Smed-Albino* (48) gene is now renamed as *Smed-foxP*. The new classification of all *Smed* Fox genes can be found in Table 1 (also in S1 Table). We did not to relate the subclassification of family genes between species, since it could cause a misleading annotation (i.e. *foxD2* genes of *Smed* and *Hsap* are not directly homologs). The analysis of their protein domains shows that in addition to the FKH domain, the K family genes also contained the Forkhead associated domain (FHA) and the P family also showed a related coiled coil. Most of the proteins were enriched in nuclear localization signal (NLS) and/or nuclear export signal (NES), in accordance with their function as TFs (Fig. S6).

**Table 1.**
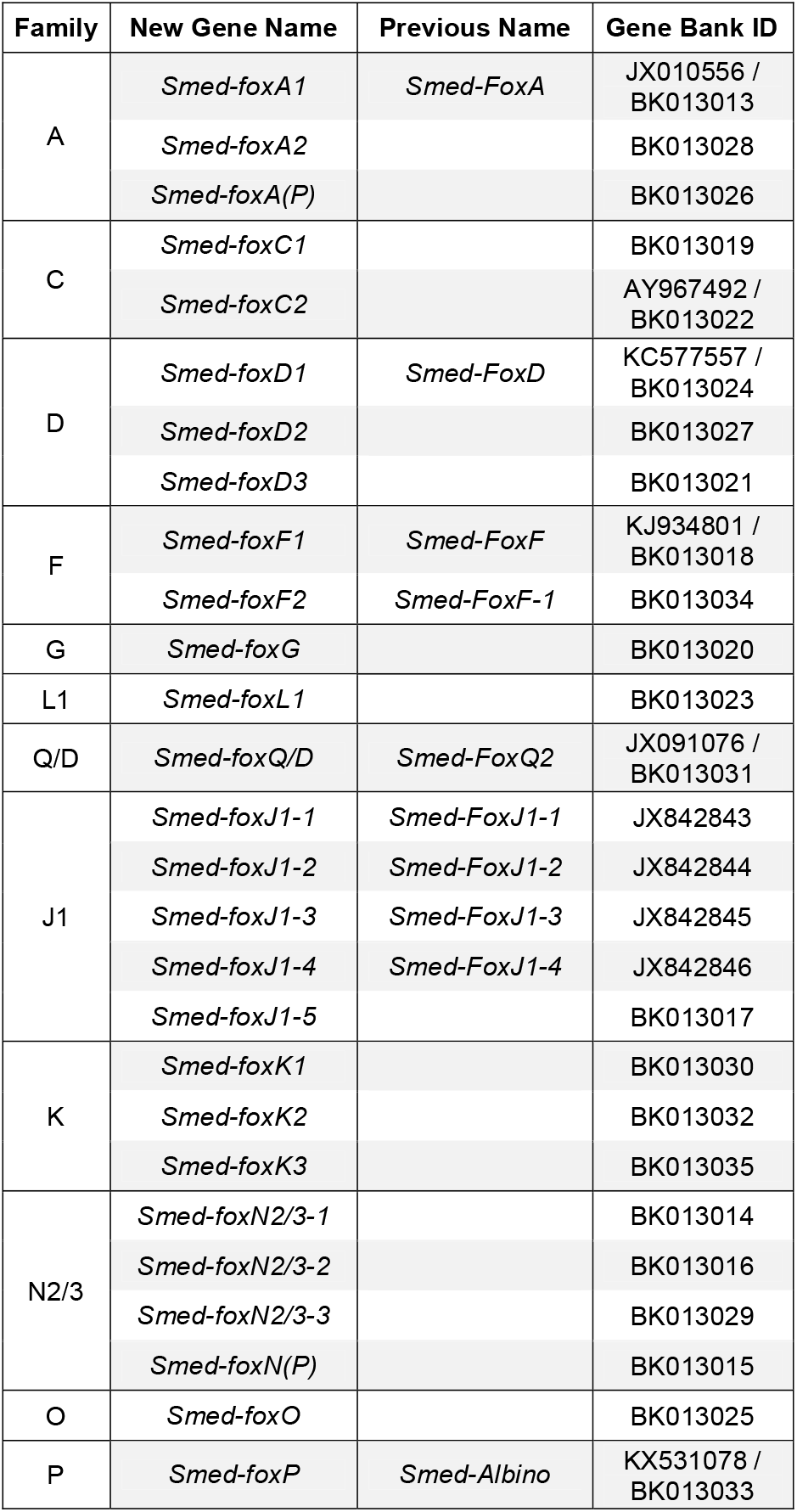
Fox genes in *Schmidtea mediterranea*. New and previous names of Fox genes in *Smed* are shown, with their corresponding GeneBank Ids.

Thanks to this new analysis, we could also confirm the loss of several Fox families in planarians (the AB, B, E, H, I, Q1, Q2, M and N1/4 families) and we could determine that most of this family losses found in planarians predate the emergence of the Platyhelminthes phylum. Besides the family loses earlier mentioned (Fig. 2), Tricladida additionally lost the J2/3 family. Interestingly, Tricladida (see turquoise lines in Fig. 4) have doubled the FKH-containing genes compared to the other Platyhelminthes, while the number of families remained constant, supporting an intrafamily diversification of Fox genes in this group (Fig. 4).

**Figure 4.**
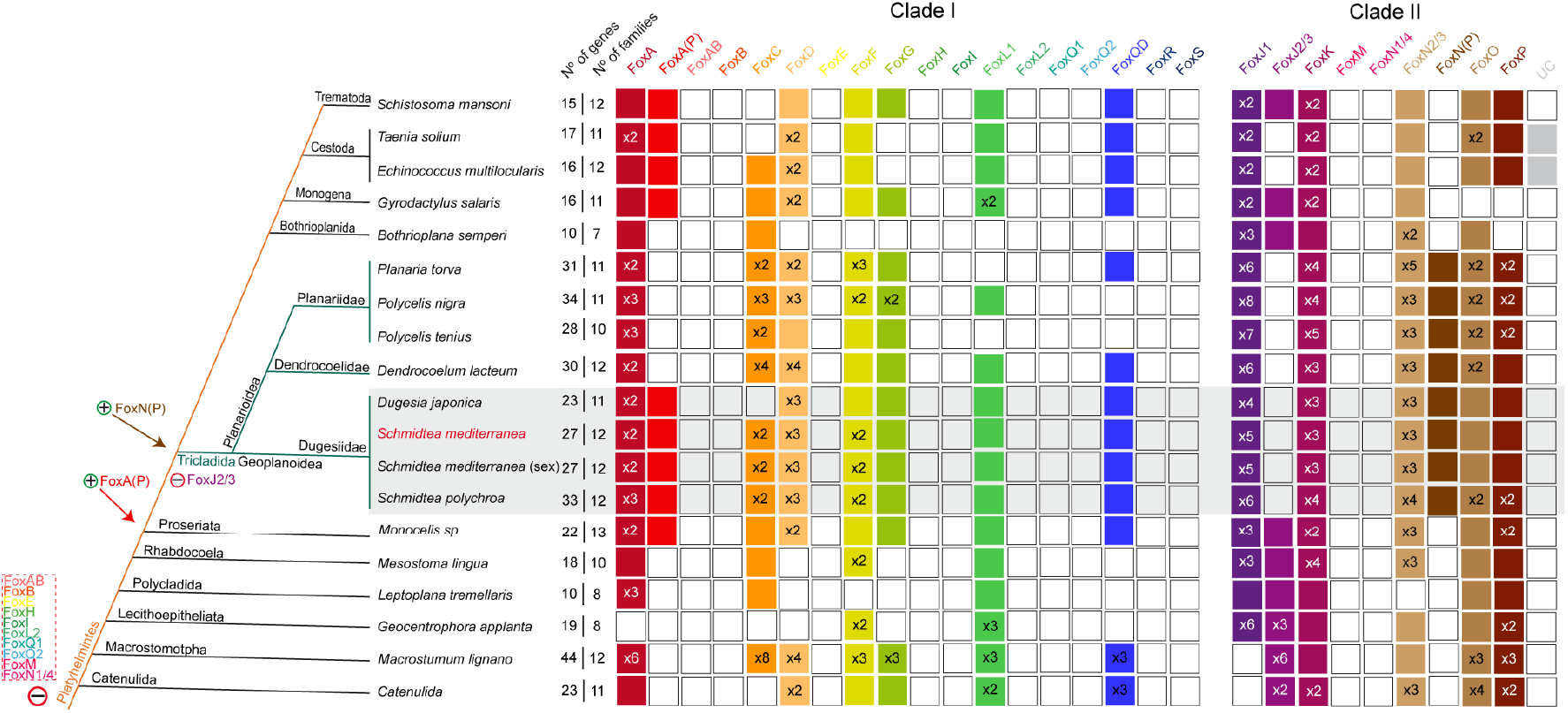
Distribution of Fox homologs reveals huge family loss and gene diversification in Platyhelminthes. Coloured boxes indicate the presence of an ortholog based on the phylogenetic analysis. When there were no evidences of ortholog, box remains white. A number (x N°) inside a box indicates paralogs per family and species. Families are divided in Clade I and II. Number of genes and number of families per species are indicated. Species were classified into Platyhelminthes and Tricladida phylogeny accordingly to (28) and (105), respectively. Light orange indicates Platyhelminthes species within turquoise lines indicate Tricladida species; grey box indicates Dugesiidae family. Species belonging to Tricladida order show different losses, gains and specialization events. Gains (+) and a losses (−) of genes are indicated. Specifically, many families were lost in Platyhelminthes (red dashed polygon). FoxA(P) and FoxN(P) origin seems to predate Proseriata and Tricladida order, respectively.

To note, the new QD family found in this study was found to be present in all Platyhelminthes, while the Q2 family is lost in all of them. To better decipher the relation between the new QD family and Q2 family, we performed a new phylogenetic analysis (Fig. 5). Increasing the number of species used outputted more confident Bayes and Bootstrap node values, supporting that a new Fox family has been uncovered, which is present in most Metazoa, including Platyhelminthes. Some genes misannotated in other families, such as Q2, B and I were reannotated as QD (Table 2, S1 File).

**Figure 5.**
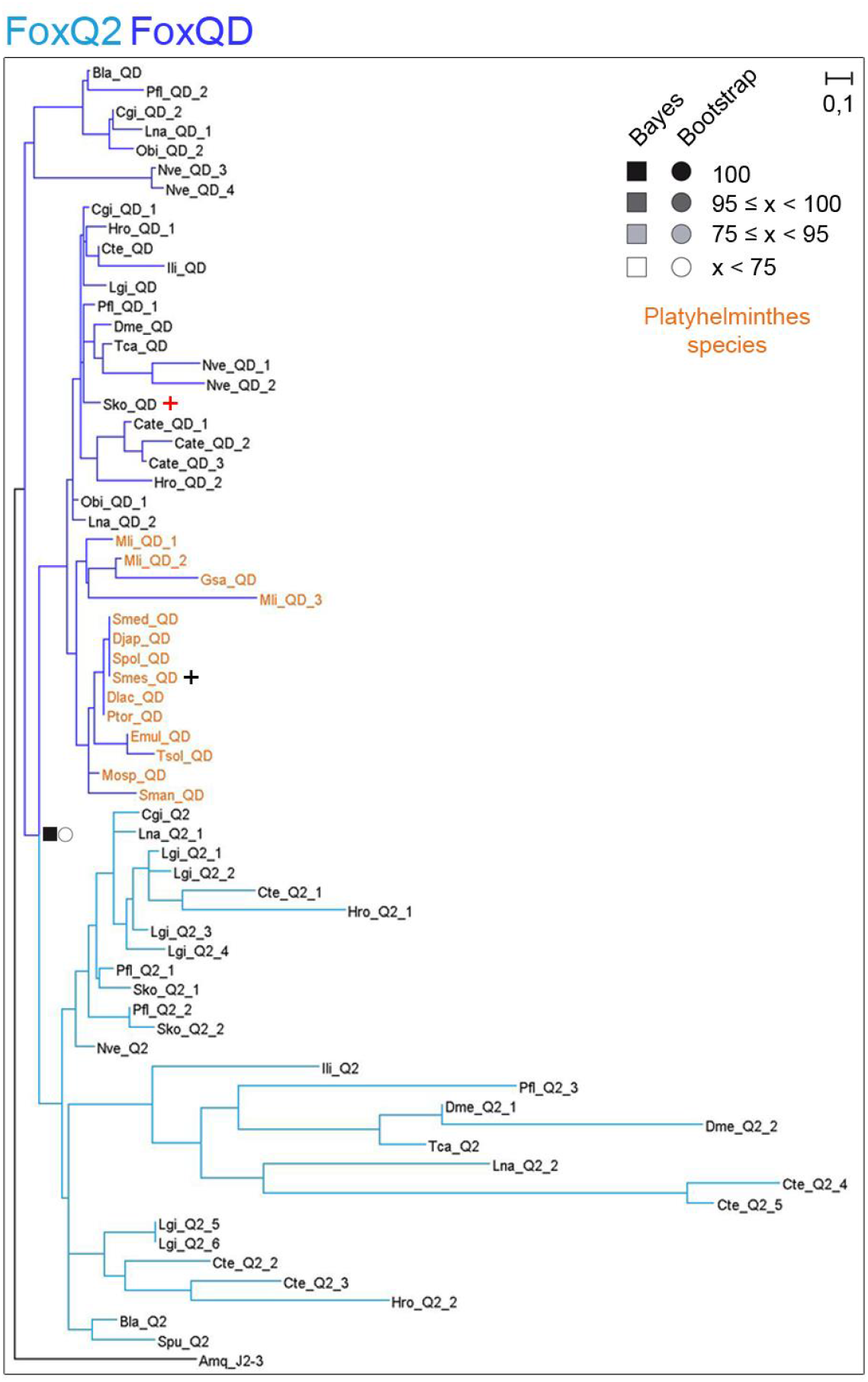
New family FoxQD is broadly found in Metazoa but missing in Vertebrata and Porifera. The ML phylogenetic trees based on FKH of Q2 and QD family evolution in all metazoan species studied in this work. At nodes, values for the approximate Bayes (square) and Likelihood (circle) ratio test are showed. Colour indicates % of confidence. An *Amq-J2/3* from the opposite clade was used as out group. Family branches are painted with the same colour as they are represented in the trees. Dark asterisk indicates new fox characterized in *Schmidtea mediterranea* (*Smed*). Red cross indicates *Saccoglossus kowalewski foxQD* gene. Platyhelminthes QD genes are coloured light orange. Aminoacidic sequences used are placed in S1 and S3 File. Scale indicates expected aminoacidic substitution per site.

**Table 2.**
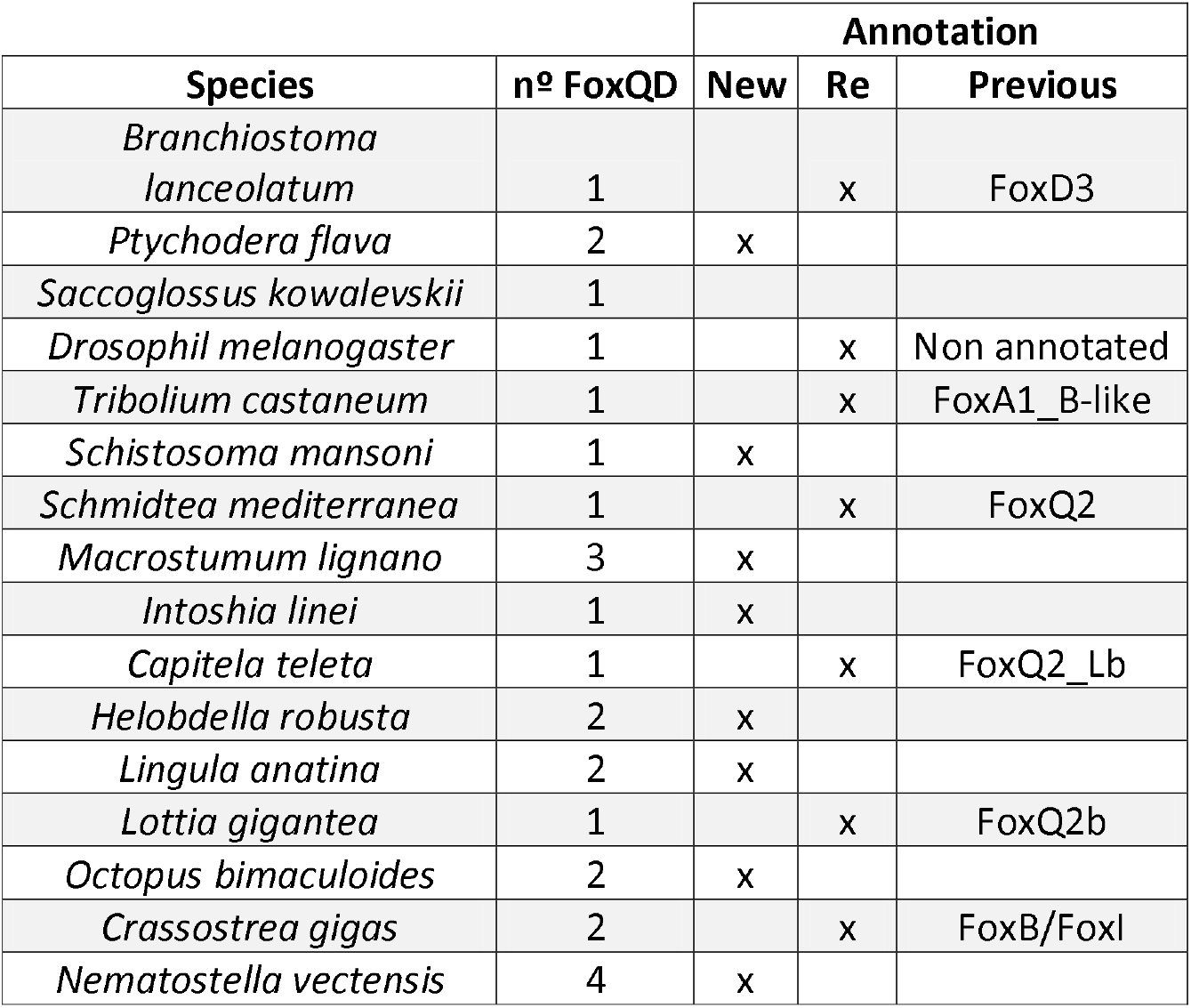
New annotation of QD genes in different metazoan species. Number of paralogs per species is indicated. They are classified as New or Re-annotated genes. In this case, the previous family is shown.

### *Schmidtea mediterranea* Fox genes appear not to be linked in the genome and to have drifted through evolution

Regarding the relative position of Fox genes in the genomes of Metazoa, we can see how some of them typically present a linkage such as in the case of the families C-F-L1-Q1 or D-E (26,49). When comparing their genomic position relative to other coding genes, Irimia et al (50) demonstrated that some of the Fox genes had retained microsynteny across metazoans with a variety of genes. Although the *Smed* genome is not assembled at chromosome level, we examined the genomic neighbourhood of *Smed* Fox genes. The only genes present in the same scaffold were *foxD2* and *foxA1* with ~187Kb of distance between them (S1 Table). Although the distance is less than the 0.1% of genome size (49) there are no other reports of A-D family linkage, meaning that no canonical Fox genes linkages are found in this assembly version of *Smed* genome. In order to verify if, despite this, there was some level of microsynteny conservation, we took an orthology-based approach similar to the one used to identify orthologous lincRNAs between distant species (51) using humans as a comparison, as well as manually checking the existence of already described microsyntenic pairs. In both cases, we found no conserved microsynteny (S5 File). Additionally, we decided to perform whole-genome alignments between the different scaffolds to examine the inter-paralog syntenic relationships (Fig. 6, S7). However, the synteny seems to be broken in all cases with most of the alignments falling exclusively into repeating elements such as LINEs.

**Figure 6.**
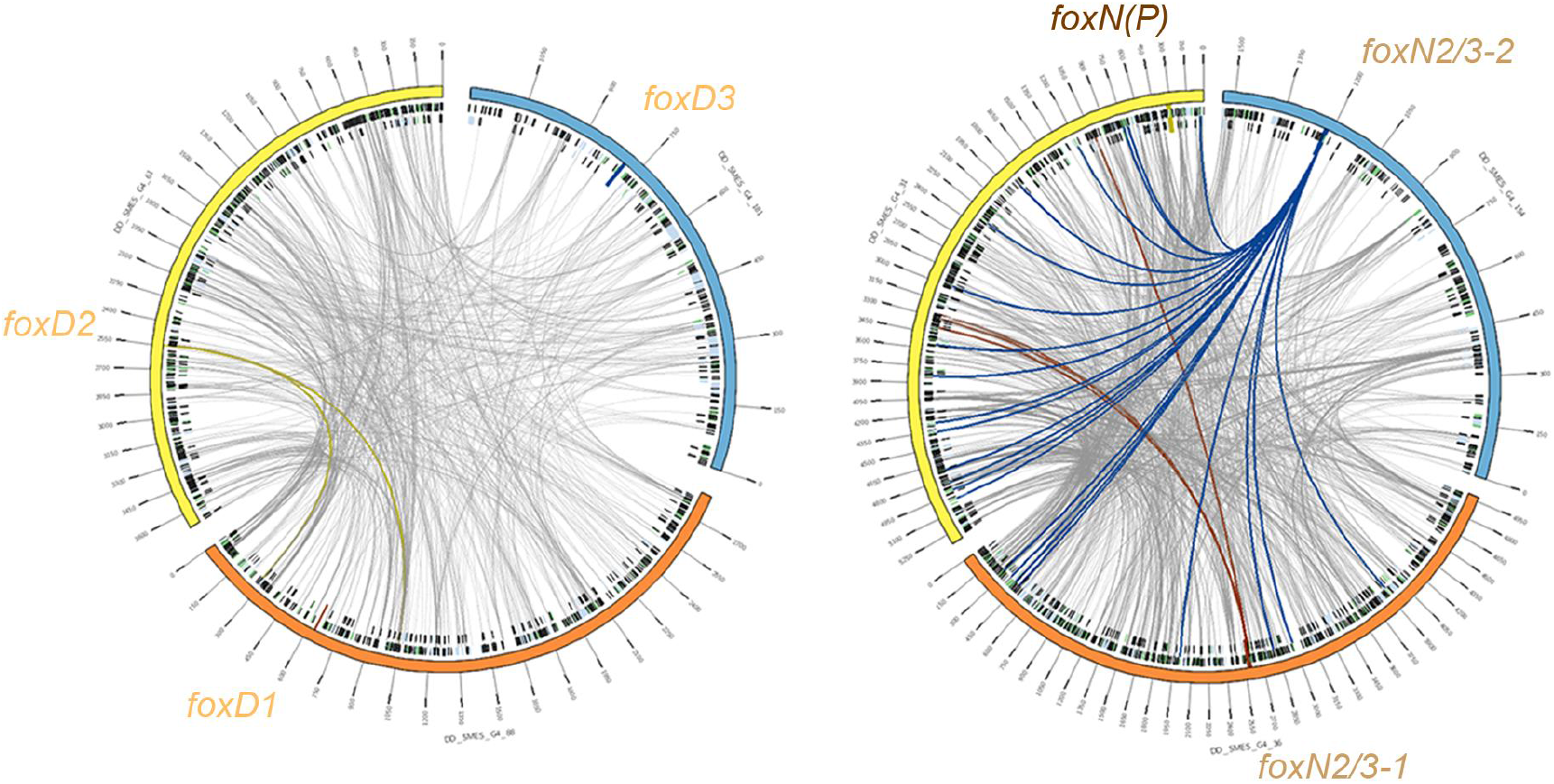
Fox paralogs do not present syntenic relationships in *Smed* genome. Alignments between scaffolds containing same-family Fox genes are represented with Circos. The Circos representation is composed of two tracks: In the outer ring the scaffolds containing Fox genes are labelled with their name (each tick representing 150kb); in the inner ring, the repeating elements (29) coloured in green (LINEs), blue (TLR) and black (simple repeats and other). Repeats are filtered to be shown only when greater than 1kb. Grey lines connecting the scaffolds are the representation of the alignments, filtered to be shown only when greater than 1Kb. In each scaffold, the region corresponding to the Fox gene (+-5Kb) is represented as a perpendicular darker region, and all the links that fall onto it are coloured accordingly.

Our data suggest that the Fox families found to be linked in other species (C-F-L1-Q1 and D-E) (26,49) are not linked in *Smed*, and *Smed* Fox genes show no microsynteny with described genes. However, this analysis should be repeated when the genome of *Smed* is completely assembled.

### *Smed-foxO* controls cell death in planarians

The expression pattern and function of most *Smed* Fox genes remain unstudied. We performed an exhaustive analysis of their expression pattern by *in situ* hybridization and by an *in-silico* search in the SCSeq databases (52,53). The results show a tissue-specific pattern of expression for most of the genes which could give clues about their function (Fig. S8-9-10, S1 Table). For instance, both uncharacterised C family genes were expressed around the pharynx, which correspond to the muscle pharynx, according to the SCSeq databases (Fig. S8-9, S1 Table); the K and the N2/3 family genes were expressed ubiquitously and in some neural populations according to the SCSeq databases (Fig. S8-9-10, S1 Table). We found particularly interesting the discovery of a unique copy of *foxO*, which is the homolog of *daf-16* in *C. elegans*, known for its role in increasing longevity (54). Nowadays *foxO* is known to be crucial in controlling metabolism and oxidative stress and also participates in the regulation of genes related to developmental processes, such as cell death, DNA repair and cell cycle (41). In good nutrient conditions, AKT phosphorylates three specific sites in Smed-FoxO (55,56), which leads to its ubiquitin degradation thanks to the 14-3-3 domains. Smed-FoxO conserves those three domains (Fig. S9a), suggesting that it is regulated by AKT. *Smed-foxO* was found to be expressed ubiquitously and according to the SCSeq databases it was present in specific cell types, such as neoblasts, neurons, parenchymal, secretory and epidermal cells (Fig. S8-9-10, S1 Table). To gain insight into its role in planarians, we inhibited its function by RNAi in starving animals. qPCR demonstrates *Smed-foxO* mRNA downregulation in *Smed-foxO* RNAi animals (Fig. S9b). After two weeks of inhibition, half of knockdown animals presented unpigmented zones distributed across their body (Fig. 7a). Analysis of the CNS (anti-synapsin) and the pharynx (DAPI) demonstrates that those structures are smaller and not properly maintained in *Smed-foxO* RNAi animals (Fig. 7b). The problems in tissue turnover could arise by unpaired cell proliferation and/or cell death. To test these possibilities, we analysed M phase cells through PH3 staining, and apoptosis by TUNEL and caspase-3 assay. Interestingly, cell proliferation was unaffected after two weeks of inhibition (Fig. S9c). However, both TUNEL and caspase-3 assays demonstrate that the apoptotic response that normally takes place in starved planarians, which are shrinking, was inhibited in *Smed-foxO* RNAi animals (Fig. 7c, 7d).

**Figure 7.**
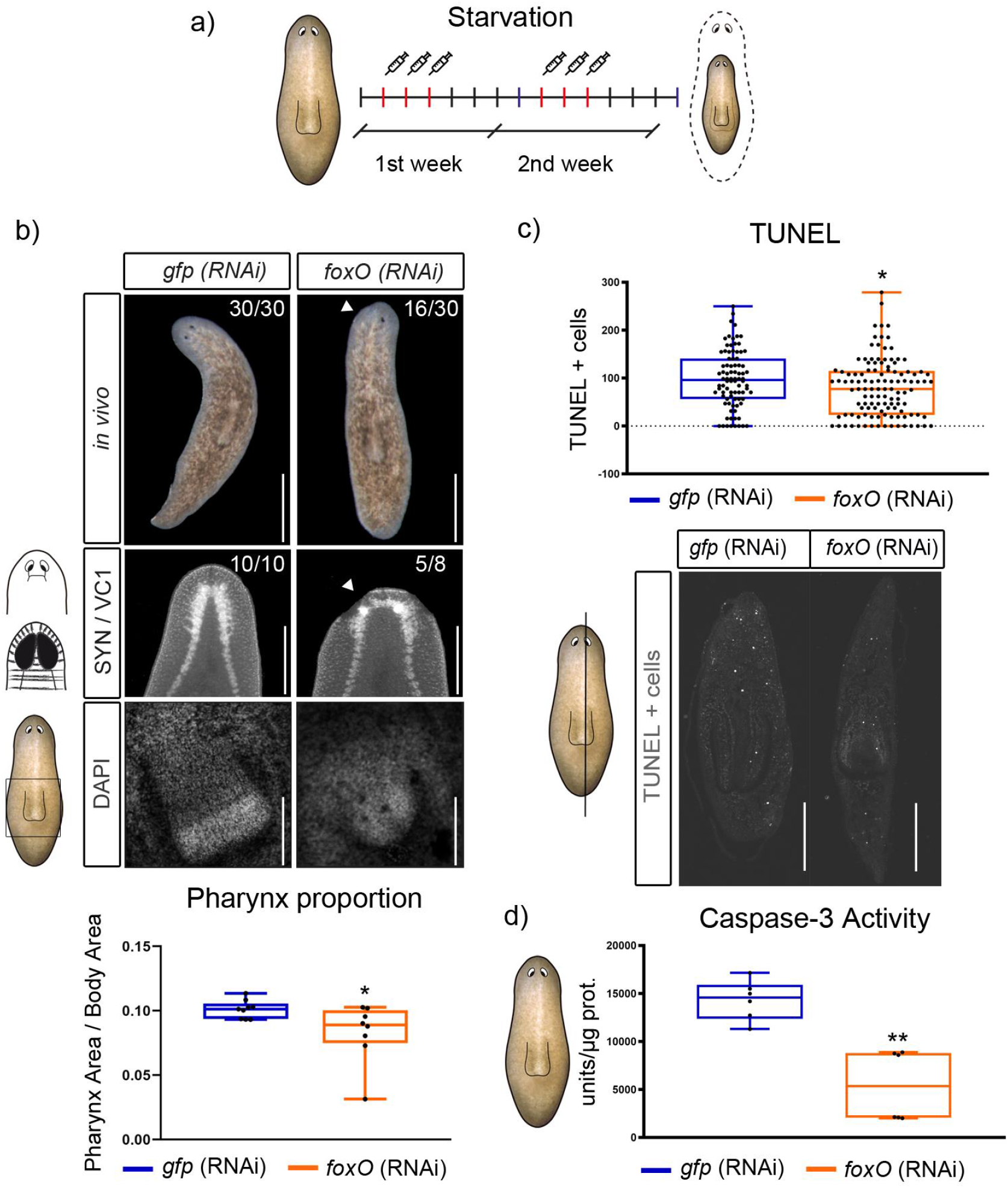
Starving *foxO* (RNAi) animals show tissue disruption and a reduction of cell death. **a** Schematic depicting RNAi procedure. **b** 50% of *in vivo foxO* knockdown animals presented unpigmented regions (white arrow), neural tissue disappearance (white arrow in synapsin immunostained animals, and pharynx size reduction, as shown by DAPI staining and relative size quantification (controls, n>9; RNAi, n=8; \**P*<0.05). **c** Quantification of TUNEL+ cells on transversal sections show a reduction of positive cells in *foxO* (RNAi) animals compared to controls (controls, n=86; RNAi, n=117; \**P*<0.05). **d** Quantification of caspase-3 activity in *foxO* (RNAi) animals and controls shows cell death reduction (controls, n>6; RNAi, n>6; \*\**P*<0.01). Scale bars: b up and mid = 100 μm and down = 10 μm; c = 100 μm.

Overall, these results demonstrate that *Smed-foxO* is necessary for planarian cell turnover through the control of cell death but not cell proliferation.

## Discussion

### The Fox genes in Metazoa: a story of early gains and specific losses

The Fox genes evolution has been a field of interest since their discovery. Although quite rich in different families, it seems that the ancestral Fox gene was remarkably similar to the J1 family. This family has been proposed as the original Fox family, giving rise to the rest of families by gene expansion and duplication (42). Few Fox genes have been identified in choanoflagellates and in sponges (6,49) and, as seen in Fig. 2, an expansion of the families took place before the origin of cnidarians, when most of the Clade II families and some Clade I families appeared. At the base of bilaterians, new families appeared: F, H and L1 (exclusive of bilaterians); R and S (vertebrate and mammal specific, respectively). This would mean that some of the Fox families are easily lost through evolution, as seems to be the case for H, I, and Q1 families. In agreement with other analysis (43,57), those families would have appeared early on the metazoans, and then were lost in a species-specific manner several times through evolution (Fig. 2). In addition, our findings support that some clades suffered massive losses, such as Ecdysozoa which has lost seven families, as it has been proposed by Mazes et al. (14), although the losses in this clade could be less if more organisms where used in the analyses. Nonetheless, the clade with the greatest number of specific losses seems to be Platyhelminthes with ten families lost (Fig. 4). This is in line with other studies that have analysed the evolution of different gene families in Platyhelminthes, such as the Wnt family, which demonstrated a great number of family losses in this clade (58–60).

### Uncovering the QD family

In this study we identified a new Fox gene family not previously described, the QD, to which some genes misannotated in other families belong to, such as Q2, B and I (Table 2, S1 File). This family originated from a paralog duplication that gave rise to both the already known Q2 family and the new QD family. According to our data, the origin of the QD family could be prior to the eumetazoan radiation although to confirm this, analysis on placozoans and ctenophores would need to be carried out. The QD family is present in almost all taxa studied and is completely lost only in vertebrates (Fig. 2).

Not only the phylogenetic data but also their different expression patterns support that QD and Q2 are different families. While Fox Q2 genes show an evolutionary conserved role in anterior brain development (44,61), *Sko foxQD* shows a completely different spatiotemporal expression (12). *Smed foxQD*, which was annotated as *foxQ2*, is not expressed in nor has a role in anterior patterning (47,62,63). We hope that, much like in the case of the uncovering of the AB family by Yu et al. (13), this new family can help to elucidate some of the inconsistencies in annotation such as in the case of *Smed* and will also contribute to a better understanding of the function of the different Fox families.

### Planarian-specific Fox: FoxA(P) & FoxN(P)

When analysing the Platyhelminthes tree, we were able to properly classify the outgrouping L1/I *Smed* Fox into the L1 family (Fig. 3b). Furthermore, having more Platyhelminthes in the phylogeny resulted in two families grouping into a new branch (Fig.3a, 3b, S4, S5). These were named *foxA(P)* and *foxN(P)* as they appeared to be divergent members of the A and N families. The L1 family also appeared to be divergent in all Tricladida and is the cause of FoxL1/I appearing as an outgroup in the metazoan tree (Fig. 1c). In contrast with the QD family, which was present all along the metazoan tree, these two new subfamilies were only present in the Platyhelminthes superphylum. We have considered them to be Platyhelminthes-specific subfamilies as they still cluster together with their main branch, but do not mix with the other members. In the case of the N(P), it appears to be Triclad-specific. However, a more exhaustive phylogenetic analysis should be performed to support this observation. We propose that the huge losses that took place before the origin of this clade may have caused the duplication and divergence that ultimately lead to the formation of the A(P) and N(P) subfamilies. *foxN(P)* expression resembles the other N family genes suggesting that they may have a similar role in regulating SNC, but further functional experiments should be performed to test it. Resemblance between Fox genes or other TFs expression patterns might suggest the co-option of the function of other Fox families (12) or other TF families (64,65).

### Tricladida may have suffered a genome reorganisation

Regardless of the extensive above-mentioned variations in family numbers due to gains and losses, the number of Fox genes present across organisms seems to be constant. The conserved number is roughly 23 genes with some obvious exceptions such as humans or sponges (Fig. 2). This regularity may seem surprising, but as Fernandez and Gabaldon (66) pointed out, family losses are usually compensated by gene duplications of the remaining family members (as appears to be the case in Platyhelminthes). We cannot discard the possibility that family losses were compensated by the emergence of *de novo* genes, which could cluster in pre-existing families due to artefacts of the clustering methods (60). Another interesting aspect to examine is the relation between the number of Fox genes and the number of Fox families. When we study this connection in Platyhelminthes we find the opposite; the number of families remains constant while the number of Fox genes increases in Tricladida (Fig. 4). This could be explained in three ways: i) several tandem duplications occurred before the branching of Tricladida, ii) a partial genome duplication event happened and affected the Fox genes, iii) a whole genome duplication (WGD) event happened. Currently, we do not have enough data to support any of these hypotheses. Other studies that have found gene duplications in Platyhelminthes species propose that WGD events could have occurred (58,59,67,68). In *Smed*, we found every Fox gene in a different scaffold (except for *foxA1* and *foxD1*) and we could not find any trace of microsynteny retained in the Fox gene regions, not even within the *Smed* genome comparing paralogs. Thus, in the *Smed* genome we cannot observe any indicator of WGD although a massive reorganization, erasing any trace of it, could have occurred. For clarification, we would need a number of Tricladida available genomes of sufficient quality. Regrettably, the *Smed* genome is the only high quality Tricladida genome currently accessible.

### Cell death regulation by *Smed-foxO* is conserved in planarians

Several Fox families have been lost or have been expanded in Platyhelminthes. Interestingly, the FoxO family is one of the few that shows a unique family member. FoxO has crucial roles in controlling molecular process related with aging and cancer (41, 69). FoxO senses oxidative stress and responds through regulating cell motility, stress resistance, or cell death (70). FoxO also has a conserved role in maintaining cellular energy homeostasis by coordinating cellular supplies and demands (39). When nutrients are available, InsulinR is activated and FoxO is phosphorylated by AKT, which inhibits its entrance into the nucleus (55,56,71). Under conditions of growth factor limitation or other stresses, FoxO enters the nucleus and inhibits mTORC1. Our data shows that *Smed-foxO* is probably regulated by AKT, since it conserves the phosphorylation domains and, furthermore, *Smed-foxO* RNAi inhibition impairs the apoptotic response, which is the opposite phenotype described after AKT RNAi in planarians (77). Thus, it could be proposed that in starved planarians, the limitation of nutrients inhibits the insulin pathway and AKT, allowing for the increase in levels of unphosphorylated *Smed-foxO* that can enter the nucleus. In the nucleus, *Smed-foxO* activates the apoptotic response required in starved planarians to trigger degrowth (72–77). In *Smed-foxO* RNAi animals this transcriptional activity cannot take place and neither can the apoptotic response. The reduction of cell death after *foxO* inhibition appears to be evolutionarily conserved as it has been also observed in *Hydra* (78), *C.elegans* (79,80), *Drosophila melanogaster* (20,81) and in various mammal tissue types (82–85).

*Smed-foxO* could also be regulated by other pathways as the Sirtuin family, which senses cellular metabolic state and acetylates FoxO (activation). Recently, Ziman et al. have demonstrated in planarians that upon starvation and after *sirtuin-1* inhibition (*foxO* inhibition) animals display a reduction in cell death (76).

The cellular REDOX state is not only essential for cellular homeostasis but it is necessary to activate the regenerative response in several models, as well as in planarians (86). In this study we have not clarified whether *Smed-FoxO* also senses ROS levels in intact or regenerating planarians, but this will be an interesting direction for the future.

## Conclusions

As we acquire more information on the presence of the TF Fox family across metazoan species, it becomes clearer that some Fox genes were originated at the base of metazoans followed by different events of gene loss and diversification as proposed by (42,66,87). Following this thread, as the number of annotated Fox genes increase, our ability to classify them also improves up until the point where ideally no Fox would be misannotated. In the past, these errors in the annotation led to a misunderstanding of the evolution of conserved functions in different Fox families.

In this study, the new annotation allowed for the proposal of a new family present in most metazoans, the FoxQD, as well as phylum-specific families exclusively found in Platyhelminthes. The appearance of phylum-specific families might not be unique to Platyhelminthes and could have happened several times throughout evolution. To prove this theory, there is a need of a better Fox gene annotation from all across the metazoan species. Besides, the proper phylogeny of these genes is not the only benefit. Having better annotated Fox genes in different key species will also help us to understand how different gene regulatory networks and developmental processes could have evolved.

*Schmidtea mediterranea* is a unique model that raises developmental questions in an evolutionary context due of its position in lophotrochozoans, an under-studied clade, and due to its stem cell-based plasticity. The identification of 27 genes divided into 12 families will give us the bases from which to understand how the TF take part in the regulation of key molecular pathways that control major developmental roles. In particular, in this study we proved that *Smed-foxO*, which in contrast to other families is constantly present in metazoans, is evolutionary conserved to regulate cell death.

Finally, the identification of complete gene families in *Smed* will also help to understand the evolution of planarians and Platyhelminthes. Here we have seen how in the order Tricladida the number of Fox genes increased while the number of families was retained. However, we could not find traces of neither a genome (whole or partial) duplication event nor tandem duplications of the Fox genes. This indicates that in planarian ancestors a genomic reorganisation could have occurred. A larger amount of Platyhelminthes and Tricladida genomes are needed to clarify these evolutionary scenarios.

## Methods

All methods were carried out in accordance with relevant guidelines and regulations.

Experiments were performed with planarians, flatworms that do not require specific approvals.

### Sequence and phylogenetic analyses

For generating the phylogenetic trees, we first obtained the FOX protein sequences from several different sources. In some of the cases we were able to collect them from the public databases, like in the case of *Hsa* or *Xtr* (S2 Table). For the rest of the organisms a manual annotation was required. If the only resource available was a transcriptome, like in the case of *Tsol*, we used Transdecoder (v5.5.0) to obtain the translated proteins. Using HMMER with default parameters and a cutoff e-value of 1e-4 (v3.1b2) and the Pfam (88) motive of the Forkhead domain (PF00250.13), we extracted the Forkhead-containing proteins.

The whole set of translated proteins was aligned again using MAFFT (89) with the L-INS-i strategy and the aligning Forkhead domain was selected. This alignment was the input used for IQ-TREE (90) to generate the phylogenetic tree. The options used to run the webserver of IQ-TREE were the ones by default, including the automatic substitution model selector and the ultrafast bootstrap analysis, except for the number of bootstrap alignments (set at 2500), the single branch test number of replicates (set at 2000) and the approximate Bayes test option (selected). The trees were visualized using Dendroscope3 v3.6.3 (91) with the default parameters.

For *Smed* FOX domains disposition architecture the NCBI web server was used (http://www.ncbi.nlm.nih.gov/Structure/cdd/wrpsb.cgi) to identify FKH, FHA and FOXP coiled coil; (92) and (93) were used to identify NLS and NES, respectively.

### Paralog analysis

The homology relationships between *Smed* and human was assessed with the best hit in a two-way blast (v2.6.0) search against human RefSeq transcripts. For the analysis of *Smed* scaffolds, a blast of the whole genome against the whole genome with the parameters “-evalue 1e-20-word_size 100” was performed, and then the data was visualized using Circos (v0.69-9), adding a track for repeating elements. Both, the links and the repeats were filtered for only rendering those greater than 1Kb.

### Animal maintenance

Asexual clonal strain of *Smed* BCN-10 biotype were maintained in PAM water (94) as previously described (95). To keep planarian population, animals were fed twice per week with liver, and starved for a week before being used in any experiment

### Isolation of Fox genes and quantitative real-time PCR

In any experiment, TRIzol reagent (Invitrogen) was used to extract total RNA from intact planarians, and cDNA was synthesized as previously described in (77). Fox genes PCR fragments were cloned in into pGEM-T Easy (Promega) vector for dsRNA synthesis or pCRII (Life Technologies) vector to ssRNA synthesis. Nucleotide sequence data reported are available in the Third Party Annotation Section of the DDBJ/ENA/GenBank databases under the accession numbers TPA: BK010973-BK010987. Real-time PCR experiments were performed using 3 biological and 3 technical replicates for each condition. Expression levels were normalized to that of the housekeeping gene ura4. All primers used in this study are shown at S6 File.

### Whole-mount *in situ* hybridization

SP6 or T7 polymerase and DIG- or FITC-modified (Roche) were used to synthesise RNA probes *in vitro*. For colorimetric whole-mount *in situ* hybridization (WISH) the previously described (96) protocol was followed. Animals were sacrificed with 5% N-acetyl-L-cysteine (NAC), fixed with 4% formaldehyde (FA), and permeabilized with Reduction Solution.

### RNAi experiments

Double strand RNA (dsRNA) was synthesised by *in vitro* transcription (Roche) as previously described (97). Injections of dsRNA (3 × 32.2 nl) were carried into the digestive system of each animal on 3 consecutive days per week. All experiments were conducted with starved animals undergoing 2 consecutive weeks of injection, without amputation.

### Immunohistochemistry staining

Whole-mount immunohistochemistry was performed as previously described (98). Animals were sacrificed with 2% HCl and fixed with 4% FA. Animals were blocked in 1% bovine serum albumin (BSA) in 1X PBSTx 0,3% (Blocking Solution) for 2 h at RT. After blocking solution for 16 h rocking at 4 °C, primary antibodies were removed, and washes were per performed for at least 4 hours. Secondary antibodies were diluted in blocking solution for 16 h rocking at 4 °C.

The following antibodies used in these experiments: mouse anti-synapsin (anti-SYNORF1, 1:50; Developmental Studies Hybridoma Bank, Iowa City, IA, USA), mouse anti-VC1 (anti-arrestin, 1:15000, kindly provided by Professor K. Watanabe), rabbit anti-phospho-histone H3 (Ser10) (D2C8) (PH3) (1:500; Cell Signaling Technology). The secondary antibody used was Alexa 488-conjugated goat anti-mouse (1:400; Molecular Probes, Waltham, MA, USA) Nuclei were stained with DAPI.

### Caspase 3 activity assay

At the end of the second week of RNAi inhibition, protein extraction was performed. BioRad protein reagent was used to obtain protein concentration of the cell lysates. Fluorometric analysis of caspase-3 activity was performed as described previously (99). Fluorescence was measured in a luminescence spectrophotometer (Perkin-Elmer LS-50) using Fluostar Optima microplate fluorescence reader (BMG Labtech), applying the following settings: excitation, 380 nm; emission, 440 nm. 20 mg of protein extract was used to determine enzyme activity, incubating for 2 hours at 37°C with 20 μM caspase-3 substrate Ac-DEVD-AMC or 2 ml from a stock of 1 mg/ml for a final volume of 150 μl. Three technical replicates were analysed per condition.

### TUNEL assay on paraffin sections

Animals were sacrificed with 2%HCl and fixed with 4%PFA. Paraffin embedding and sectioning were carried out as previously described in (100). Slides were de-waxed and re-hydrated as previously described in (101). Sections were treated as described previously in (102) and after the dewaxing step, they were incubated with Proteinase K (20μg/ml for 10 min at room temperature). Finally, the ApopTag Red In situ Apoptosis Detection Kit (CHEMICON, S7165) was used, following manufacture’s protocol.

### Imaging and quantification

WISH images were captured with a ProgRes C3 camera from Jenoptik (Jena, TH, Germany). A Zeiss LSM 880 confocal microscope (Zeiss, Oberkochen, Germany) was used to obtain confocal images of whole-mount immunostainings and TUNEL staining.

Representative confocal stacks for each experimental condition are shown. Cell counting of PH3+ and TUNEL staining was carried out by eye quantification in a previous defined area of each animal. Areas are schematically indicated in each figure. The total number of PH3+ cells was divided by the animal area. For TUNEL quantification, TUNEL positive cells were counted in at least 30 representative transversal sections per animal. The number of positive cells were divided by the mean area of the all sections in each animal. Images were blind analysed and later grouped according to each genotype. At least two animals were analysed per condition.

### Statistical analysis and presentation

Statistical and presentation analyses were performed using GraphPad Prism 8. Two-sided Student’s t-tests (α = 0.05) were performed to compare the means of 2 populations. To compare 2 populations, we used box plots depicting the median, the 25th and 75th percentiles (box), and all included data points (black dots). Whiskers extend to the largest data point within the 1.5 interquartile range of the upper quartile and to the smallest data point within the 1.5 interquartile lower ranges of the quartile.

## Acknowledgements

The authors thank all the member of Emili Saló, Teresa Adell, Francesc Cebrià and Jordi García labs for discussion of the results and their suggestions; Jordi Paps for helpful advice and discussion while writing the manuscript, and Cristina Guijarro-Clarke for sharing genomic data.

## Funding

EPC and CHU are recipients of an FPI (Formación del Profesorado Investigador) scholarship from the Spanish Ministerio de Educación y Ciencia (MEC). The funder had no role in the study design, data collection and analysis, decision to publish, or manuscript preparation. ES and TA received funding from the Ministerio de Educación y Ciencia (grant number BFU2017-83755-P and BFU2014-56055-P). ES and TA benefits from 2017SGR-1455 from ACU (Genereliat de Catalunya). ES received funding from AGAUR (Generalitat de Catalunya: grant number 2009SGR1018). JGF received funding from the Ministerio de Educación y Ciencia (grant number BFU2017-86152-P). In no case the funder had no role in study design, data collection and analysis, decision to publish, or preparation of the manuscript.

## List of abbreviations

Amq: Amphimedon queenslandica
Bla: Branchiostoma lanceolatum
Bose: Bothrioplana semperi
Cate: Catenulia
Cgi: Crassostrea gigas
Cte: Capitella teleta
Djap: Dugesia japonica
Dlac: Dendrocoelum lacteum
Dme: Drosophila melanogaster
Emul: Echinoccocus multiocularis
FHA: Forkhead Associated Domain
FKH: Forkhead domain
Fox: Forkhead Box
Geap: Geocentrophora applanta
Gsa: Gyrodactylus salaris
Hro: Helobdella robusta
Hsa: Homo sapiens
Ili: Intoshia linei
Lept: Leptoplana tremellaris
Lgi: Lottia gigantea
Lna: Lingula anatine
Meli: Mesostoma lingua
Mli: Macrostomum lignano
Mli: Macrostumum lignano Mosp Monocelis sp
NES: Nuclear Exportation Signal
NLS: Nuclear Localization Signal
Nve: Nematostella vectensis
Obi: Octopus bimaculoides
Pfl: Ptychodera flava
Pnig: Polycelis nigra
Pten: Polycelis tenius
Ptor: Planaria torva
Sdo: Suberites domuncula
Sko: Saccoglossus kowalevskii
Sman: Schistosoma mansoni
Smed: *Schmidtea mediterranea* asexual strain
Smes: *Schmidtea mediterranea* sexual strain
Spol: Schmidtea polychroa
Spu: Strongylocentrotus purpuratus
Tca: Tribolium castaneum
TF: Transcription Factor
Tsol: Taenia solium
WGD: Whole Genome Duplicatiom
Xtr: Xenopus tropicals

## Notes

### Competing Interest Statement

The authors have declared no competing interest.

