## Supplementary Figures for "Analysis of Fox genes in *Schmidtea mediterranea* reveals new families and a conserved role of *Smed-foxO* in controlling cell death"

**The PDF file includes:**

**Figs. S1 to S9**

**Other Supplementry information of this manuscript includes the following:**

**S1 File**

In this file we have compiled all the data used for generating the phylogenetic tree of metazoa. Alongside the internal identificator used in the trees, there is the original ID, the database from where the sequences were retrieved, the species, the domain sequence and the complete protein sequence. We have also added the number of families, and genes for each of the species analysed. If the source of the data was obtained from a publication, the field is filled with the reference.

**S2 File**

Raw tree of phylogenetic Fox analysis from Metazoa species, placed in Fig. 1b.

**S3 File**

In this file we have compiled all the data used for generating the phylogenetic tree of Platyhelmintes. Alongside the internal identificator used in the trees, there is the original ID, the database from where the sequences were retrieved, the species, the domain sequence and the complete protein sequence. We have also added the number of families, and genes for each of the species analysed. If the source of the data was obtained from a publication, the field is filled with the reference.

**S4 File**

Raw tree of phylogenetic Fox analysis from Plathelmintes species, placed in Fig. 3a.

**S4 File**

In this file we can find the position of every Fox gene identified in Smed along a summary of the microsyntenic features found by Irimia et al. (1) with a Fox gene involved separated by family. We have also compiled the three upstream and downstream annotated orthologs (by best hit against human; * indicates orthologies with two-way BLAST hits) for every Fox gene in *Smed*, as well as the three upstream and downstream coding genes for each Fox gene in *Homo sapiens* whose family is present in *Smed*. Finally in a separate tab is the full list of *Hsa* Fox genes with their three upstream and downstream neighbours. Orthologies are indicated by same coloured fonts.

**S5 File.** Primers used in this study

We display: name of the primer, technique were they used and the sequences forward (Fw) and reverse (Rv) 5' to 3'.

**S1 Table. Fox genes in *Schmidtea mediterranea***. The following information is shown: genome ID and its scaffold disposition, the transcriptome ID (2), new and previous names of Fox genes in *Smed*, GeneBank IDs of new and previous *Smed* Fox genes (and the reference where they were mentioned before), and the homologs identified in relative planarian species. The expression pattern obtained by WISH and from the SCseq databases is also showed. N.D. non-detected / N.S.C. No Specific Cluster

**S2 Table. Fox protein sources.** Fox protein sources. For each species, the source of their Fox genes and the database or the previous work is indicated.

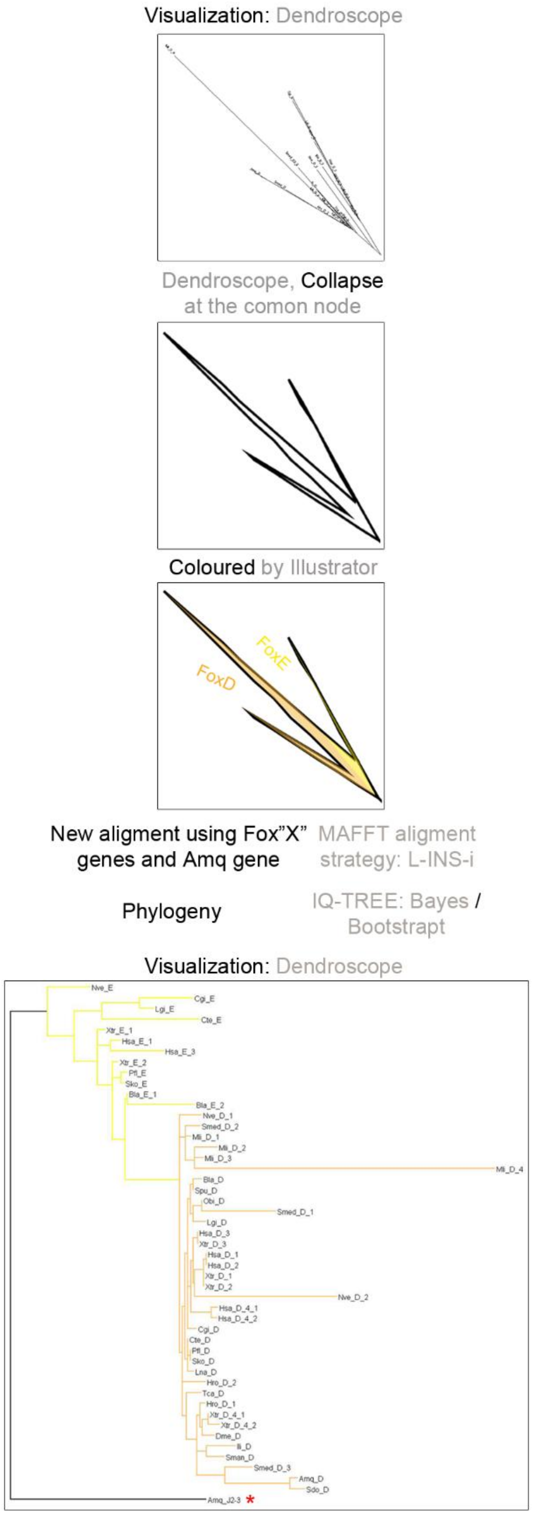


**Figure S1. Family tree representation workflow.** Families are collapse at the common node and schematic colour was added. All genes per family were collected, aligned and a new phylogenetic analysis was done. All branches belonging to the same family are coloured equal as the family represented in the main tree. To obtain a better visualization, an

*Amq* gene was used as a root (red asterisk): *Amq-foxJ2/3* for Clade I and *Amq-foxD* for Clade II genes.

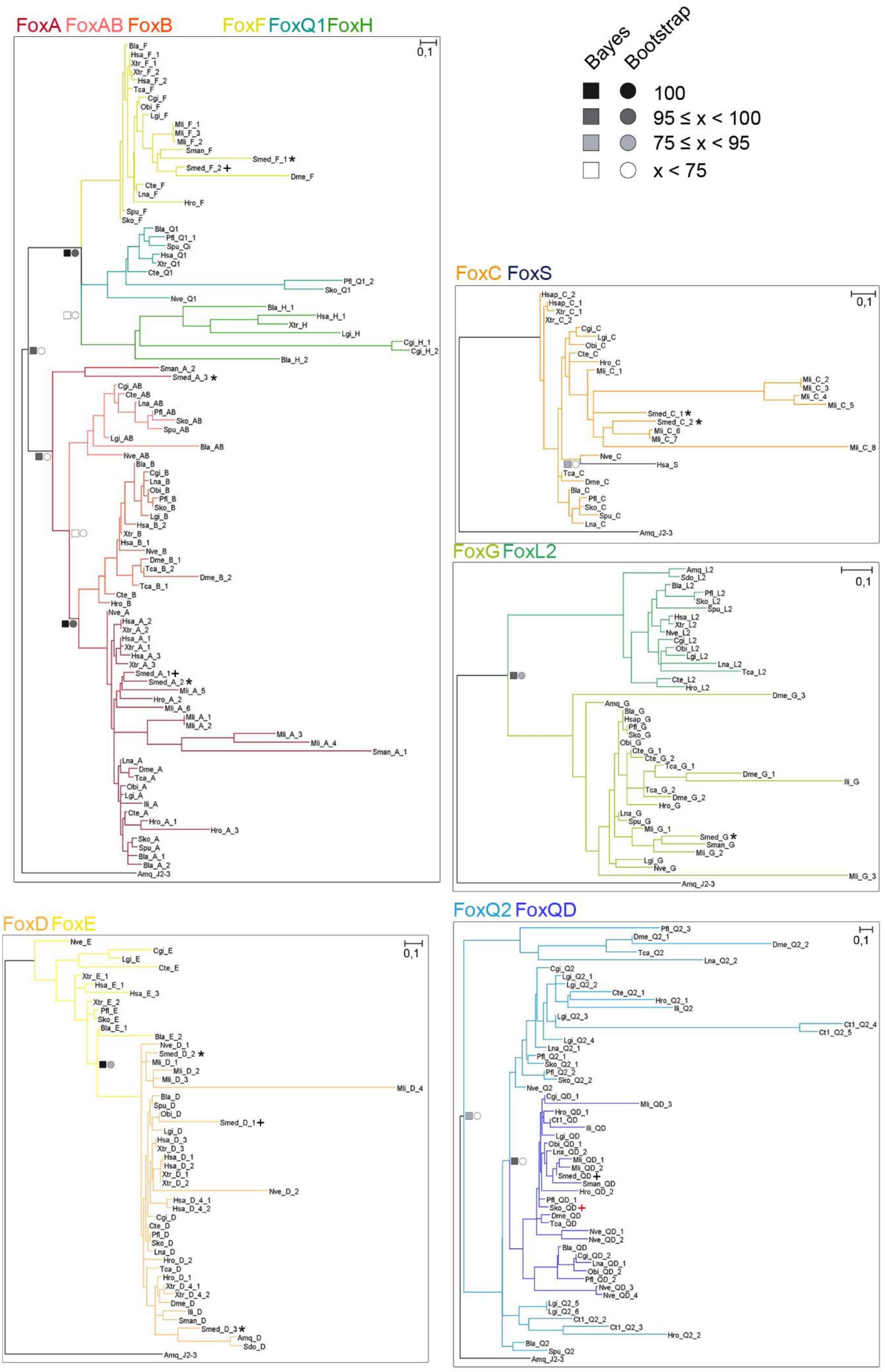
FoxA FoxAB FoxB
FoxF FoxQ1 FoxH
Bayes
Bootstrap
100
95 ≤ x < 100
75 ≤ x < 95
x < 75
FoxC FoxS
FoxG FoxL2
FoxD FoxE
FoxQ2 FoxQD
0,1

**Figure S2. Phylogenetic trees from node-sharing families.** The ML phylogenetic trees based on FKH domain. At nodes, values for the approximate Bayes (square) and Likelihood (circle) ratio test are showed. Colour indicates % of confidence. For each node-sharing families, phylogenetic trees were created using *Amq* gene from the opposite clade as out group. Family branches are painted with the same colour as they are represented in the main tree. Dark cross indicates previous characterized gene and dark asterisk indicates new fox characterized in *Schmidtea mediterranea* (*Smed*). Red cross indicates *Saccoglossus kowalewski foxQD* gene. Aminoacidic sequences used are found in Additional File 1. Scale indicates expected aminoacidic substitution per site. Species used are the following ones: *Homo sapiens* (*Hsa*), *Xenopus tropicals* (*Xtr*), *Branchiostoma lanceolatum* (*Bla*), *Strongylocentrotus purpuratus* (*Spu*), *Saccoglossus kowalewski (Sko*) and *Ptychodera flava* (*Pfl*), *Drosophila melanogaster* (*Dme*), *Tribolium castaneum* (*Tca*), *Crassostrea gigas* (*Cgi*), *Lottia gigantea* (*Lgi*), *Octopus bimaculoides* (*Obi*), *Lingula anatine* (*Lna*), *Intoshia linei* (*Ili*), *Capitella teleta* (*Cte*), *Helobdella robusta* (*Hro*), *Macrostomum lignano* (*Mli*), *Schistosoma mansoni* (*Sman*), *Nematostella vectensis* (*Nve*), *Amphimedon queenslandica* (*Amq*) and *Suberites domuncula* (*Sdo*).

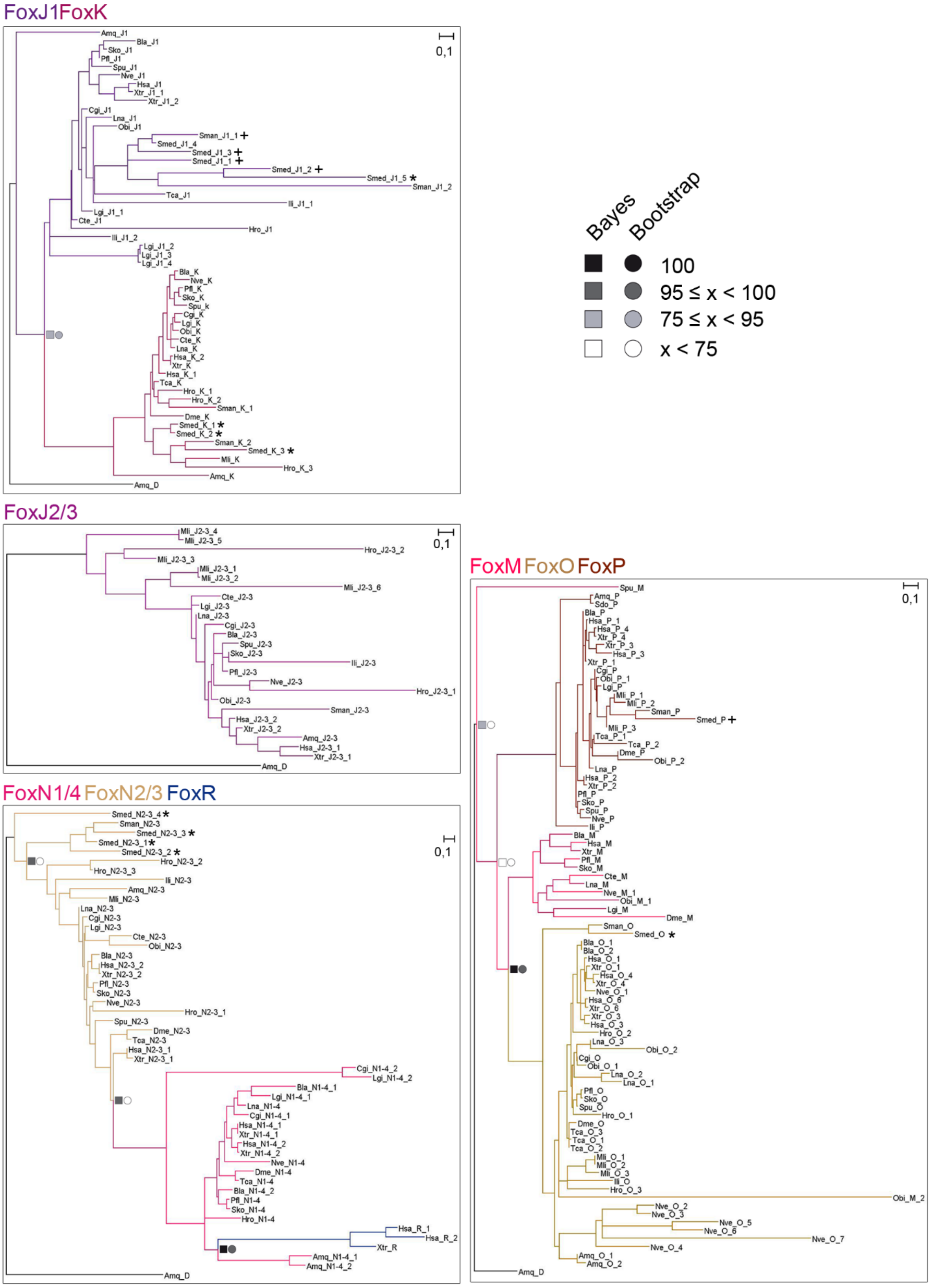


**Figure S3. Phylogenetic trees from node-sharing families.** The ML phylogenetic trees based on FKH domain. At nodes, values for the approximate Bayes (square) and Likelihood (circle) ratio test are showed. Colour indicates % of confidence. For each node-sharing families, phylogenetic trees were created using *Amq* gene from the opposite clade as out group. Family branches are painted with the same colour as they are represented in the main tree. Dark cross indicates previous characterized gene and dark asterisk indicates new fox characterized in *Schmidtea mediterranea* (*Smed*). Aminoacidic sequences used are found in Additional File 1. Scale indicates expected aminoacidic substitution per site. Species used are the following ones: *Homo sapiens* (*Hsa*), *Xenopus tropicals* (*Xtr*), *Branchiostoma lanceolatum* (*Bla*), *Strongylocentrotus purpuratus* (*Spu*), *Saccoglossus kowalewski (Sko*) and *Ptychodera flava* (*Pfl*), *Drosophila melanogaster* (*Dme*), *Tribolium castaneum* (*Tca*), *Crassostrea gigas* (*Cgi*), *Lottia gigantea* (*Lgi*), *Octopus bimaculoides* (*Obi*), *Lingula anatine* (*Lna*), *Intoshia linei* (*Ili*), *Capitella teleta* (*Cte*), *Helobdella robusta* (*Hro*), *Macrostomum lignano* (*Mli*), *Schistosoma mansoni* (*Sman*), *Nematostella vectensis* (*Nve*), *Amphimedon queenslandica* (*Amq*) and *Suberites domuncula* (*Sdo*).

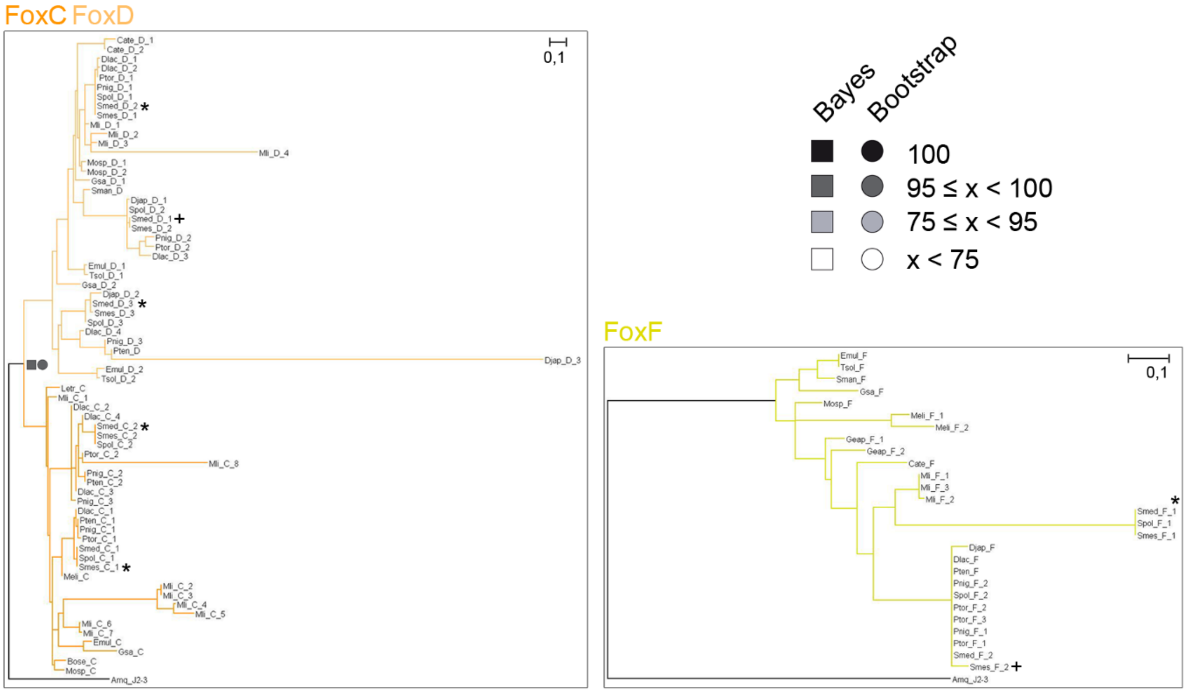


**Figure S4. Phylogenetic trees from node-sharing families.** The ML phylogenetic trees based on FKH domain. At nodes, values for the approximate Bayes (square) and Likelihood (circle) ratio test are showed. Colour indicates % of confidence. For each node-sharing families, phylogenetic trees were created using *Amq* gene from the opposite clade as out group. Family branches are painted with the same colour as they are represented in the main tree. Dark cross indicates previous characterized gene and dark asterisk indicates new fox characterized in *Schmidtea mediterranea* (*Smed*). Aminoacidic sequences used are found in Additional File 3. Scale indicates expected aminoacidic substitution per site. Species used are the following ones: *Taenia solium* (*Tsol*), *Echinoccocus multiocularis* (*Emul*), *Gyrodactylus salaris* (*Gsa*), *Bothrioplana semperi* (*Bose*), *Macrostumum lignano* (*Mli*), *Monocelis sp.*(*Mosp*), *Mesostoma lingua* (*Meli*), *Leptoplana tremellaris* (*Lept*), *Geocentrophora applanta* (*Geap*), *Catenulia* (*Cate*), *Planaria torva* (*Ptor*), *Polycelis nigra* (*Pnig*), *Polycelis tenius* (*Pten*), *Dendrocoelum lacteum* (*Dlac*), *Dugesia japonica* (*Djap*), the sexual strain of *Schmidtea mediterranea* (*Smes*) and *Schmidtea polychroa* (*Spol*).

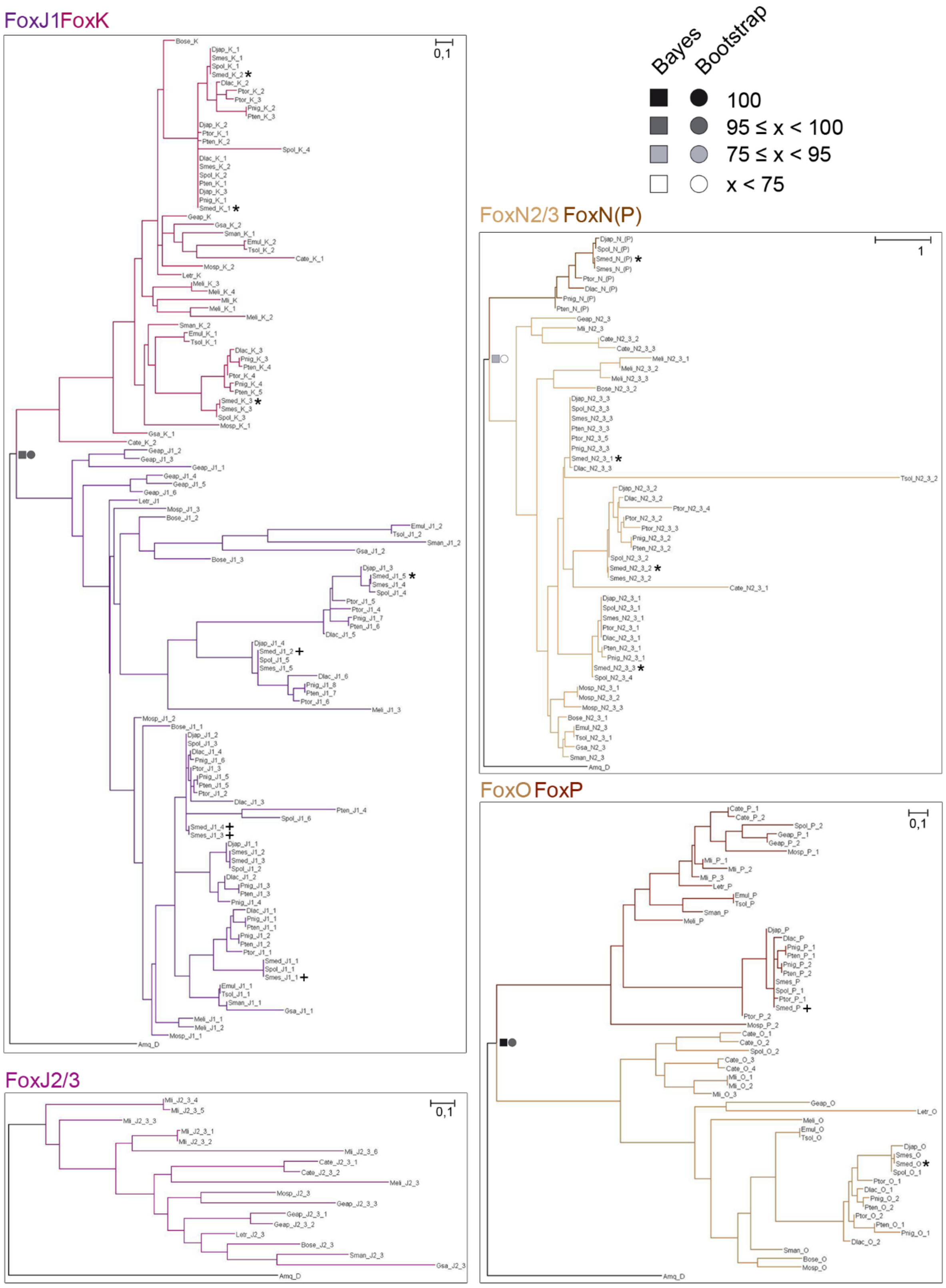


**Figure S5. Phylogenetic trees from node-sharing families.** The ML phylogenetic trees based on FKH domain. At nodes, values for the approximate Bayes (square) and Likelihood (circle) ratio test are showed. Colour indicates % of confidence. For each node-sharing families, phylogenetic trees were created using *Amq* gene from the opposite clade as out group. Family branches are painted with the same colour as they are represented in the main tree. Dark cross indicates previous characterized gene and dark asterisk indicates new fox characterized in *Schmidtea mediterranea* (*Smed*). Aminoacidic sequences used are found in Additional File 3. Scale indicates expected aminoacidic substitution per site. Species used are the following ones: *Taenia solium* (*Tsol*), *Echinoccocus multiocularis* (*Emul*), *Gyrodactylus salaris* (*Gsa*), *Bothrioplana semperi* (*Bose*), *Macrostumum lignano* (*Mli*), *Monocelis sp.*(*Mosp*), *Mesostoma lingua* (*Meli*), *Leptoplana tremellaris* (*Lept*), *Geocentrophora applanta* (*Geap*), *Catenulia* (*Cate*), *Planaria torva* (*Ptor*), *Polycelis nigra* (*Pnig*), *Polycelis tenius* (*Pten*)*, Dendrocoelum lacteum* (*Dlac*), *Dugesia japonica* (*Djap*), the sexual strain of *Schmidtea mediterranea* (*Smes*) and *Schmidtea polychroa* (*Spol*).

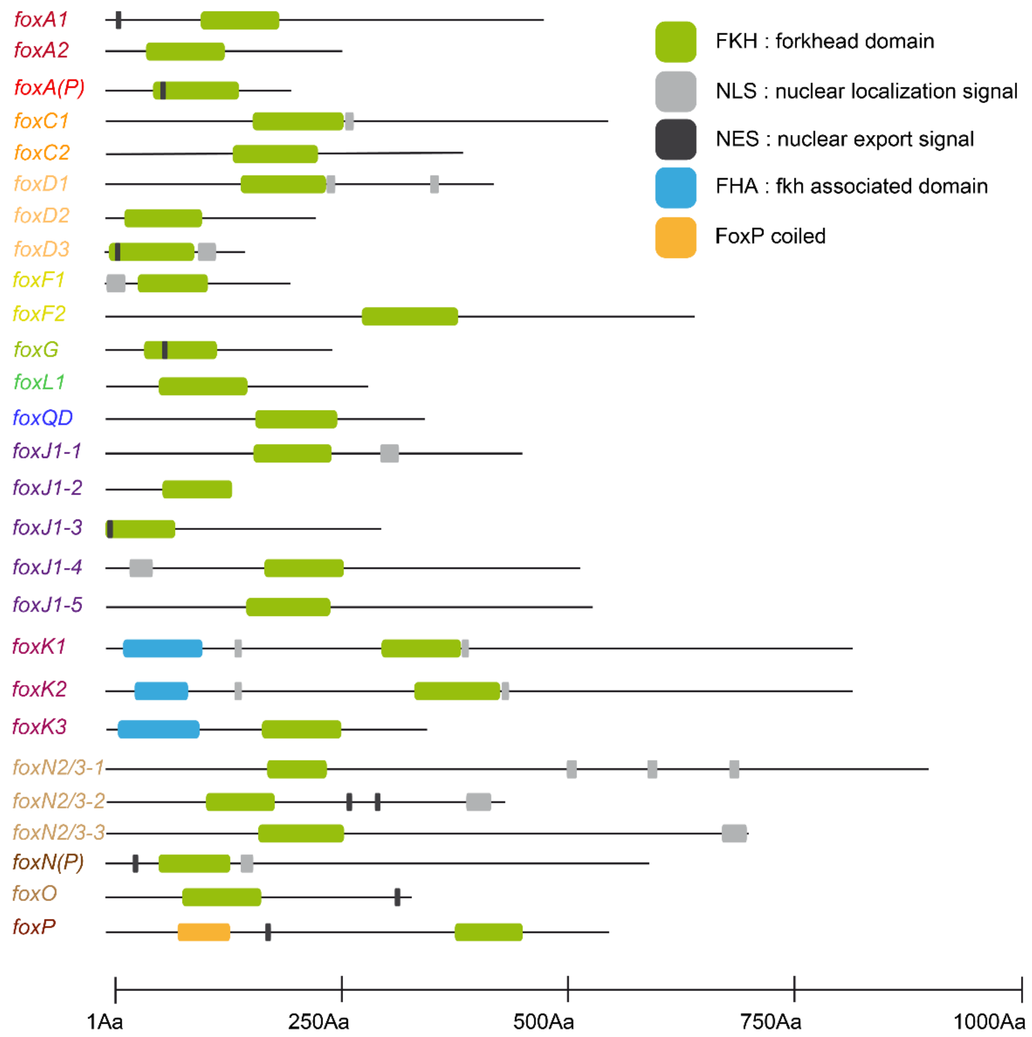


**Figure S6. Domains of Fox proteins in *Schmidtea mediterranea*.** Conserved domains in planarians: Forkhead domain (green), Forkhead associated domain (blue), FoxP coiled (yellow), nuclear localization signal (light grey) and nuclear export signal (dark grey).

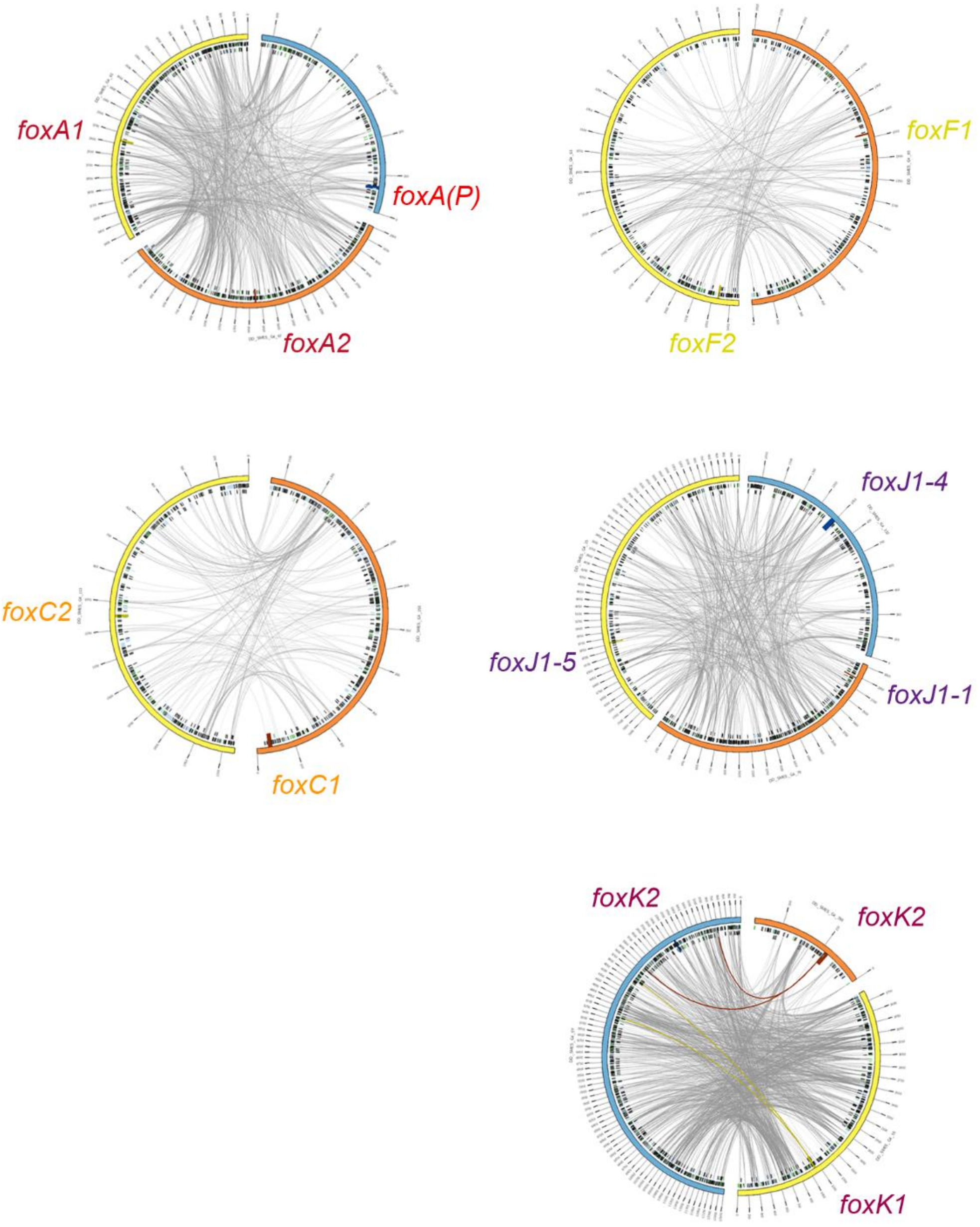


**Figure S7. *Smed* genomic alignments between scaffolds containing same-family Fox genes represented with Circos**. The Circos representation is composed of two tracks: In the outer ring, the scaffolds containing Fox genes are labelled with their name and in the outer ring (each tick representing 150kb); in the inner ring, the repeating elements (2) coloured in green (LINEs), blue (TLR) and black (simple repeats and other). Repeats are filtered to be shown only when greater than 1kb. Grey lines connecting the scaffolds are the representation of the alignments, filtered to be shown only when greater than 1Kb. In each scaffold, the region corresponding to the Fox gene (+-5Kb) is represented as a perpendicular darker region, and all the links that fall onto it are coloured accordingly.

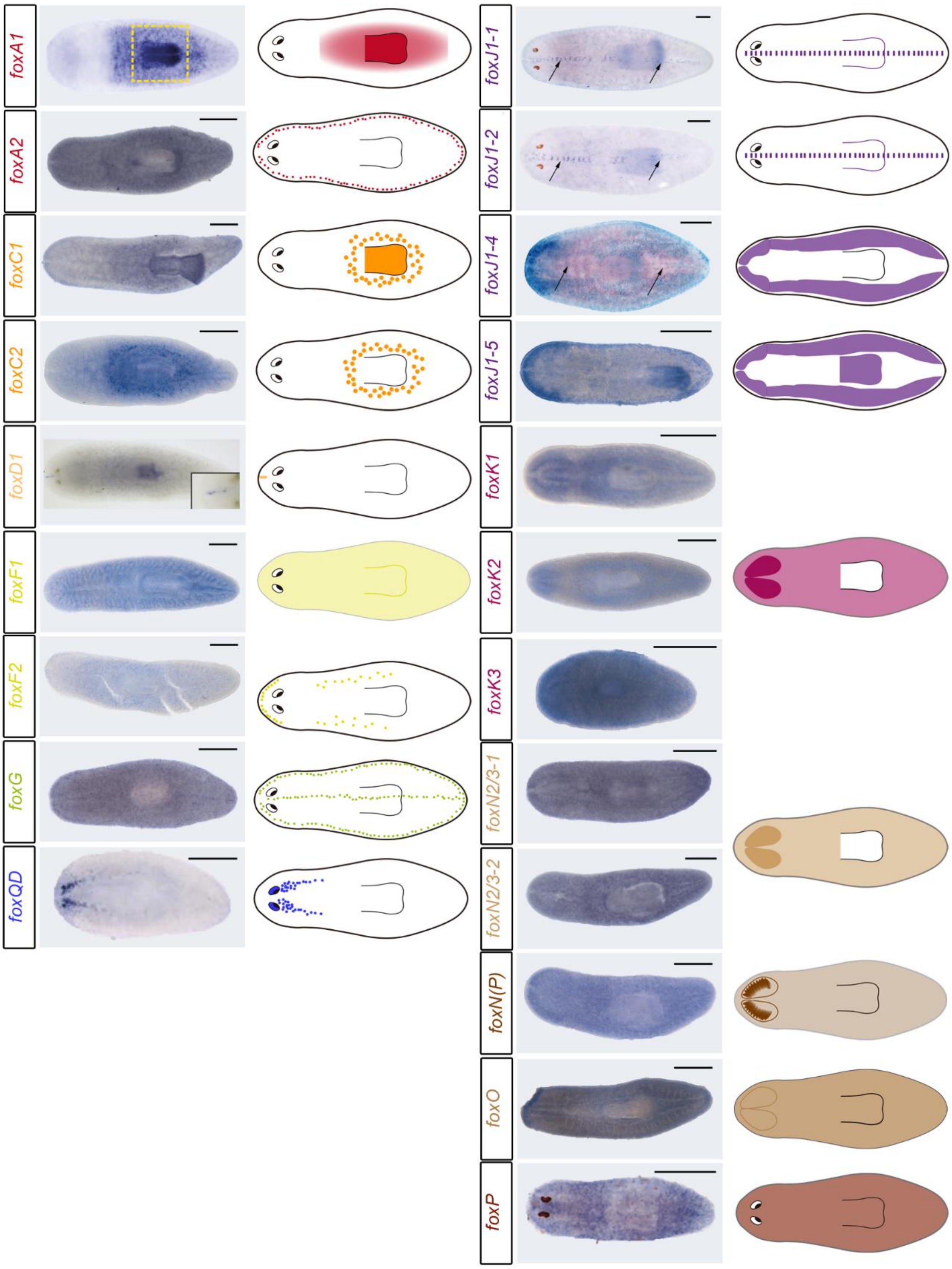


**Figure S8. *Smed-fox* genes show specific expression patterns.** WISH of Fox genes in planarians. For all genes a schematic cartoon showing expression is added. Gene names appear laterally at each image. Within the A family, *foxA1* was expressed in the pharynx (yellow dashed square) and its progenitors (3); *foxA2* was marginally expressed in a dotted pattern all along the animal body; the new *foxA(P)* could not be detected (the SCSeq data suggests that it could be expressed in early epidermal progenitors and/or non-ciliated neurons). Both *foxC* genes were expressed around the pharynx. *foxC2* was also expressed at the pharynx itself. *foxD1* was expressed in muscle cells, a magnification of the anterior tip was added (4,5). The other two D family genes were not detected by FISH. SCSeq data reveals that *foxD2* could be present in some neural progenitors and non-ciliated neuronal cell type. *foxD3* could be found at different muscle cell types, early epidermal progenitors and neurons otf+2 (Fig. S7). One member of the F family, *foxF1*, was previously described to be expressed in muscle (non-body wall) and pigment cells (6). *foxF2* was expressed in cells in the margin of the head and in the lateral dorsal part of the animal, between the pharynx and the margin of the organism. *foxG* was expressed in a subset of muscles all along the DV margin, in the dorsal midline and some scattered cells in dorsal and ventral part. *foxL1* was not detected by ISH. The SCSeq data indicates that it could be found in a muscular pharynx cell type (Fig. S7). *foxQ/D* was expressed in differentiated eye cells (rhabdomeric photoreceptor neurons), some brain progenitors and in ventral nerve cords (7). Three J1 family paralogs previously described were expressed in ciliated cells, black arrows indicate different patterns. They disposition could be more dorsally or more ventrally depending on the gene (8). The non-previously described *foxJ1-5* was also expressed in the epidermis more concentrated in the head area and in the pharynx. The three K family genes were expressed in the ubiquitously and specifically in the CNS. The N family contains 4 genes: *foxN2/3-1* and *foxN2/3-2* were expressed ubiquitously and the last one was also expressed in the SNC; *foxN2/3-3* was not detected by ISH; *foxN(P)* was expressed in the brain branches. *foxO* was expressed ubiquitously, and SCSeq reveals specific expression in some neural, epidemic and parenchymal populations. *foxP* is ubiquitously expressed (9). The expression of some Fox genes was not possible to be detected, although we designed at least two different riboprobes. Scale bars: 250 µm, as exception: *foxA1* = 10 µm, *foxD1* = 10 µm, *foxD1* = 100 µm, *foxQD* = 500 µm, *foxJ1-1* and *foxJ1-2* = 200 µm, *foxJ1-4* = 300 µm and *foxP* = 500 µm.

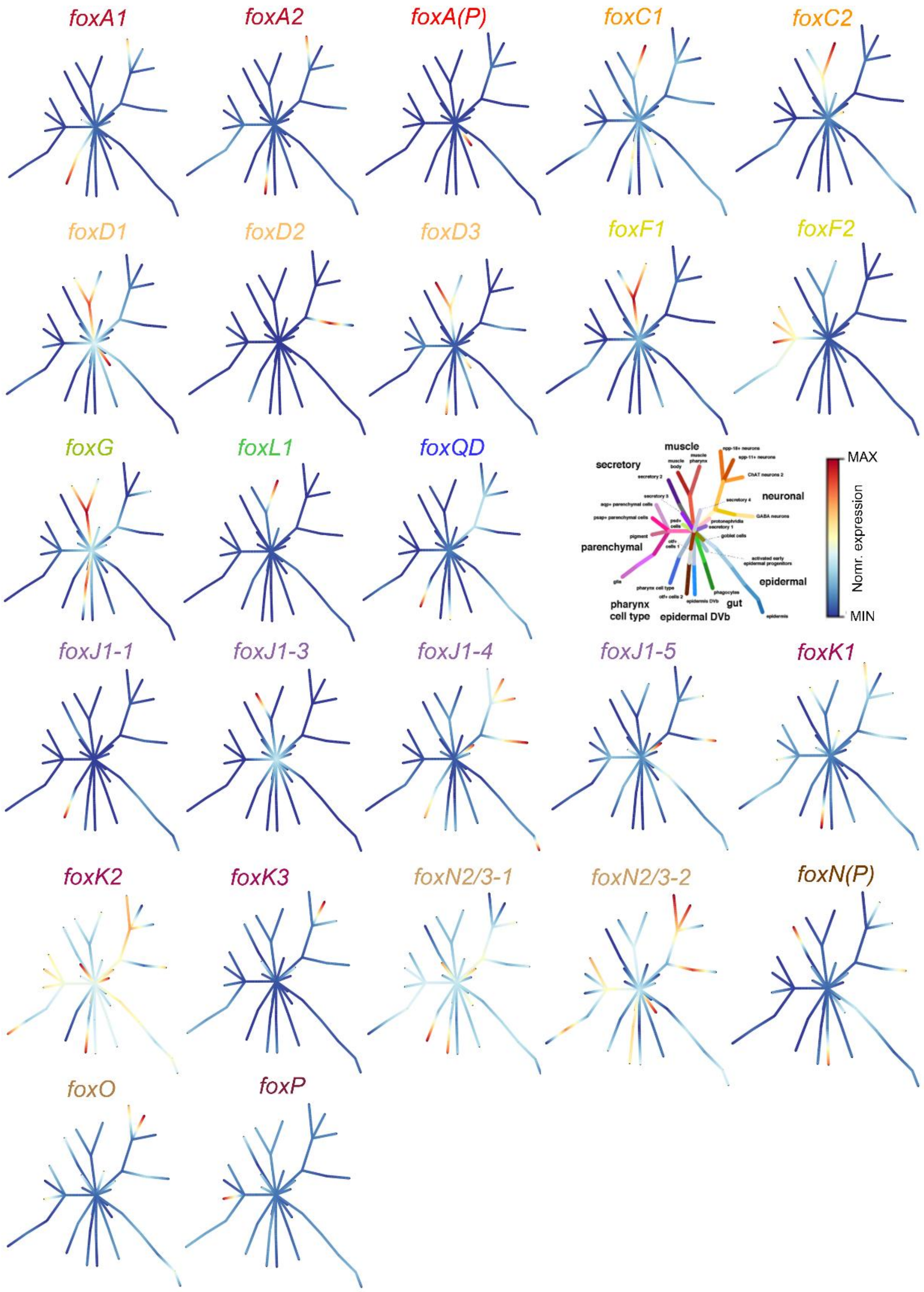


**Figure S9.** Graphical representation of *Smed-fox* genes expression during cell differentiation obtained from Plass et al. (10)

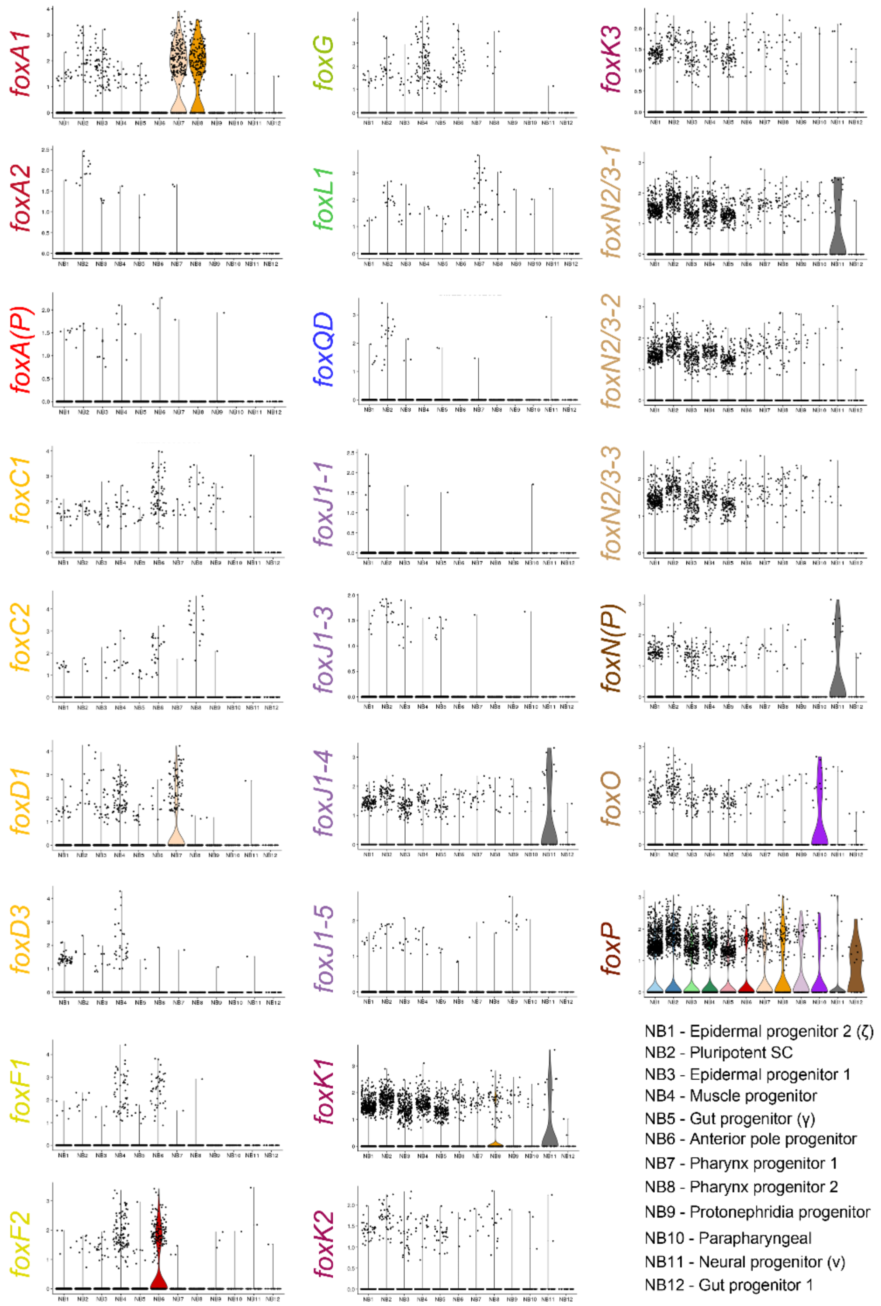


**Figure S10.** Graphical representation of *Smed-fox* genes expression in neoblasts obtained from Zeng et al. (11).

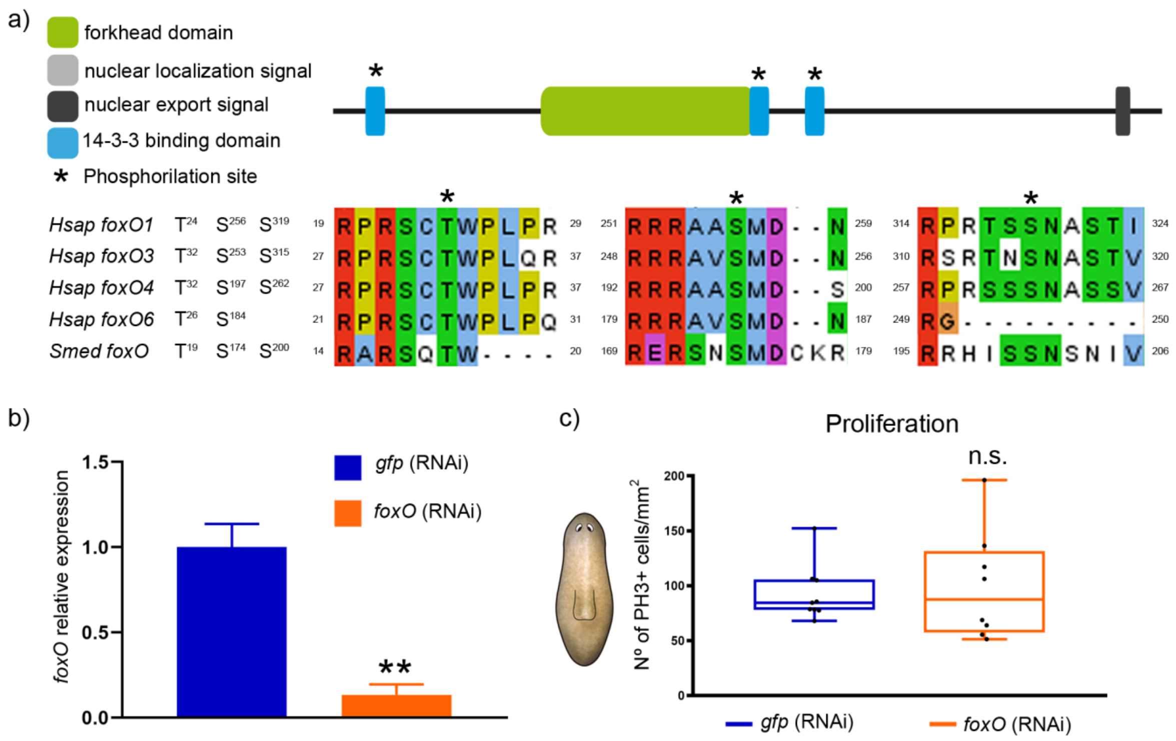


**Figure S11. *Smed-foxO* inhibition does not affect cell proliferation. a** Schematic illustration of conserved domains in Smed-FoxO. Alignment of FOXO amino acid sequences from *Homo sapiens* (*Hsap*) and *Schmidtea mediterranea* (*Smed*) showing high level of conservation of the three phosphorylation sites (*). **b** qRT-PCR analysis quantifying *Smed-foxO* expression after one week of *Smed-foxO* inhibition, proving that it was down-regulated. Relative expression is plotted as $2^{-\Delta\Delta CT}$ values. Data are plotted as mean and error bars are s. d. (***P*<0.01). **c** Quantification of PH3+ cells shows no differences between control and *Smed-foxO* (RNAi) animals after two weeks of inhibition (controls, n>7; RNAi, n>7; n.s.). In the schematic drawing the square indicates the region analysed.
